# Exploring Diverse Binding Mechanisms of Broadly Neutralizing Antibodies S309, S304, CYFN-1006 and VIR-7229 Targeting SARS-CoV-2 Spike Omicron Variants: Integrative Computational Modeling Reveals Balance of Evolutionary and Dynamic Adaptability in Shaping Molecular Determinants of Immune Escape

**DOI:** 10.1101/2025.04.15.649027

**Authors:** Mohammed Alshahrani, Vedant Parikh, Brandon Foley, Gennady Verkhivker

**Author notes:** (M.A); (V.P.); (B.F.); (G.V). Correspondence; (G.V).

## Abstract

Evolution of SARS-CoV-2 has led to the emergence of variants with increased immune evasion capabilities, posing significant challenges to antibody-based therapeutics and vaccines. The cross-neutralization activity of antibodies against Omicron variants is governed by a complex and delicate interplay of multiple energetic factors and interaction contributions. In this study, we conducted a comprehensive analysis of the interactions between the receptor-binding domain (RBD) of the SARS-CoV-2 spike protein and four neutralizing antibodies S309, S304, CYFN1006, and VIR-7229. Using integrative computational modeling that combined all-atom molecular dynamics (MD) simulations, mutational scanning, and MM-GBSA binding free energy calculations, we elucidated the structural, energetic, and dynamic determinants of antibody binding. Our findings reveal distinct dynamic binding mechanisms and evolutionary adaptation driving broad neutralization effect of these antibodies. We show that S309 targets conserved residues near the ACE2 interface, leveraging synergistic van der Waals and electrostatic interactions, while S304 focuses on fewer but sensitive residues, making it more susceptible to escape mutations. The analysis of CYFN-1006.1 and CYFN-1006.2 antibody binding highlights broad epitope coverage with critical anchors at T345, K440, and T346, enhancing its efficacy against variants carrying the K356T mutation which caused escape from S309 binding. Our analysis of broadly potent VIR-7229 antibody binding to XBB.1.5 and EG.5 Omicron variants emphasized a large and structurally complex epitope, demonstrating certain adaptability and compensatory effects to F456L and L455S mutations. Mutational profiling identified key residues crucial for antibody binding, including T345, P337, and R346 for S309, and T385 and K386 for S304, underscoring their roles as evolutionary "weak spots" that balance viral fitness and immune evasion. The results of this energetic analysis demonstrate a good agreement between the predicted binding hotspots and critical mutations with respect to the latest experiments on average antibody escape scores. The results of this study dissect distinct energetic mechanisms of binding and importance of targeting conserved residues and diverse epitopes to counteract viral resistance. Broad-spectrum antibodies CYFN1006 and VIR-7229 maintain efficacy across multiple variants and achieve neutralization by targeting convergent evolution hotspots while enabling tolerance to mutations in these positions through structural adaptability and compensatory interactions at the binding interface. The results of this study underscore the diversity of binding mechanisms employed by different antibodies and molecular basis for high affinity and excellent neutralization activity of the latest generation of antibodies.

## Introduction

The emergence of the severe acute respiratory syndrome coronavirus 2 (SARS-CoV-2) has spurred extensive research into understanding its structure, infection mechanisms, and immune responses. The SARS-CoV-2 Spike (S) glycoprotein is central to viral transmission and immune evasion, characterized by remarkable conformational flexibility [1-15]. The S1 subunit of S protein includes the N-terminal domain (NTD), receptor-binding domain (RBD), and conserved subdomains SD1 and SD2. The NTD facilitates initial host cell attachment, while the RBD binds to the angiotensin-converting enzyme 2 (ACE2) receptor, transitioning between "up" and "down" conformations to modulate receptor and antibody accessibility [1-15]. SD1 and SD2 stabilize the prefusion state and orchestrate membrane fusion, highlighting the S protein’s adaptability and complexity [10-18]. Biophysical studies have revealed the thermodynamic and kinetic principles governing its functional transitions, emphasizing mechanisms that balance receptor binding, membrane fusion, and immune escape [16-18]. The extensive array of cryo-electron microscopy (cryo-EM) and X-ray structures of S protein variants of concern (VOCs) in various functional states, along with their interactions with antibodies has provided significant insights into the virus’s adaptability [19-25]. A critical component of the immune response to SARS-CoV-2 is the production of antibodies that target various regions of the S protein, which plays a central role in viral entry into host cells. High-throughput yeast display screening and deep mutational scanning (DMS) have revolutionized our understanding of the escape mutation profiles associated with the RBD residues of the SARS-CoV-2 spike protein. These advanced techniques have enabled researchers to systematically map the functional epitopes targeted by human anti-RBD neutralizing antibodies leading to a comprehensive classification of these Antibodies into distinct epitope groups (A–F) [26]. The recent study expanded this approach to characterize the epitope distribution of antibodies elicited by post-vaccination BA.1 infection and identified the mutational escape profiles for 1,640 RBD-binding Abs, which were classified into 12 epitope groups [27]. In this classification, groups A–C consist of antibodies targeting the ACE2-binding motif that are highly effective at blocking the interaction between the virus and its host receptor, making them crucial for neutralization. However, their efficacy is often compromised by mutations in key residues within the ACE2-binding site, such as K417, E484, and N501, which are frequently observed in variants of concern. Group D antibodies such as REGN-10987, LY-CoV1404, and COV2-2130, bind to the epitope 440–449 on the RBD and are further divided into D1 and D2 subgroups. Groups E and F are subdivided into E1–E3 and F1–F3, respectively, covering the front and back of the RBD. Groups E and F represent antibodies that target regions outside the ACE2-binding motif, with each group being further subdivided into three subgroups (E1–E3 and F1–F3). These subgroups collectively cover both the "front" and "back" surfaces of the RBD, providing a comprehensive view of the antigenic landscape. Notably, groups E and F correspond to class 3 and class 4 antibodies in earlier classification systems [28], highlighting their role in recognizing non-overlapping epitopes that contribute to a diverse immune response. Group E antibodies are particularly sensitive to mutations at residues G339, T345, and R346, which are located near the N-terminal region of the RBD. These residues play a critical role in stabilizing the RBD structure and facilitating interactions with certain antibodies. In contrast, group F antibodies are affected by mutations at residues F374, T376, and K378, as well as, in some cases, V503 and G504.

S309 (group E) and S304 antibodies (group F1) represent two distinct classes of neutralizing antibodies (Figure 1). Despite their shared ability to neutralize the virus, these antibodies exhibit significant differences in their binding epitopes, mechanisms of action, and neutralization profiles. S309 targets a conserved epitope near the N343 glycosylation site within the RBD (Figure 1A). This region is adjacent to but does not overlap with the ACE2-binding motif [29,30]. S309 does not directly compete with ACE2 for binding. Instead, it induces conformational changes in the RBD that stabilize the "down" conformation, reducing the availability of the "up" state required for ACE2 engagement [29,30]. The epitope recognized by S309 is highly conserved across SARS-CoV-2 variants (Figure 1A) and related sarbecoviruses, such as SARS-CoV-1, making it effective against a broad range of coronaviruses [31]. S309 induces subtle conformational changes in the RBD that disrupt its interaction with ACE2. The cryo-EM structures of the BQ.1.1, XBB.1 and BN.1 RBDs bound to the S309 antibody and human ACE2 explain the preservation of antibody binding through conformational selection, altered ACE2 recognition and immune evasion [32]. The structures demonstrated that S309 binds to both BQ.1.1 and XBB.1 RBDs [32] in a binding pose is indistinguishable from that observed when it is bound to the Wu-WT [29,30] or the BA.1 RBD variant [33]. The S371F mutation, which is present in BA.2, BA.5, BQ.1.1, XBB.1, XBB.1.5 leads to conformational changes of the RBD helix comprising residues 364–372 that are sterically incompatible with the glycan N343 conformation observed in S309-bound S structures. Together with residue N440, position R346 is a part of the epitope of S309 and important for binding S309 (Figure 1A). R346S alone was not sufficient to alter S309 binding, but R346S in combination with P337L enhanced resistance to S309 [34]. R346K in VOC Omicron BA.1.1 reduced sensitivity to S309 [35]. S309 can synergize with other antibodies targeting distinct epitopes, such as those within the ACE2-binding motif, to create a multi-layered defense against viral escape mutations [36]. Due to its conserved epitope and non-competitive mechanism, S309 demonstrates robust neutralization activity against variants of concern (VOCs), including Alpha, Beta, Gamma, Delta, and Omicron [37,38].

**Figure 1.**
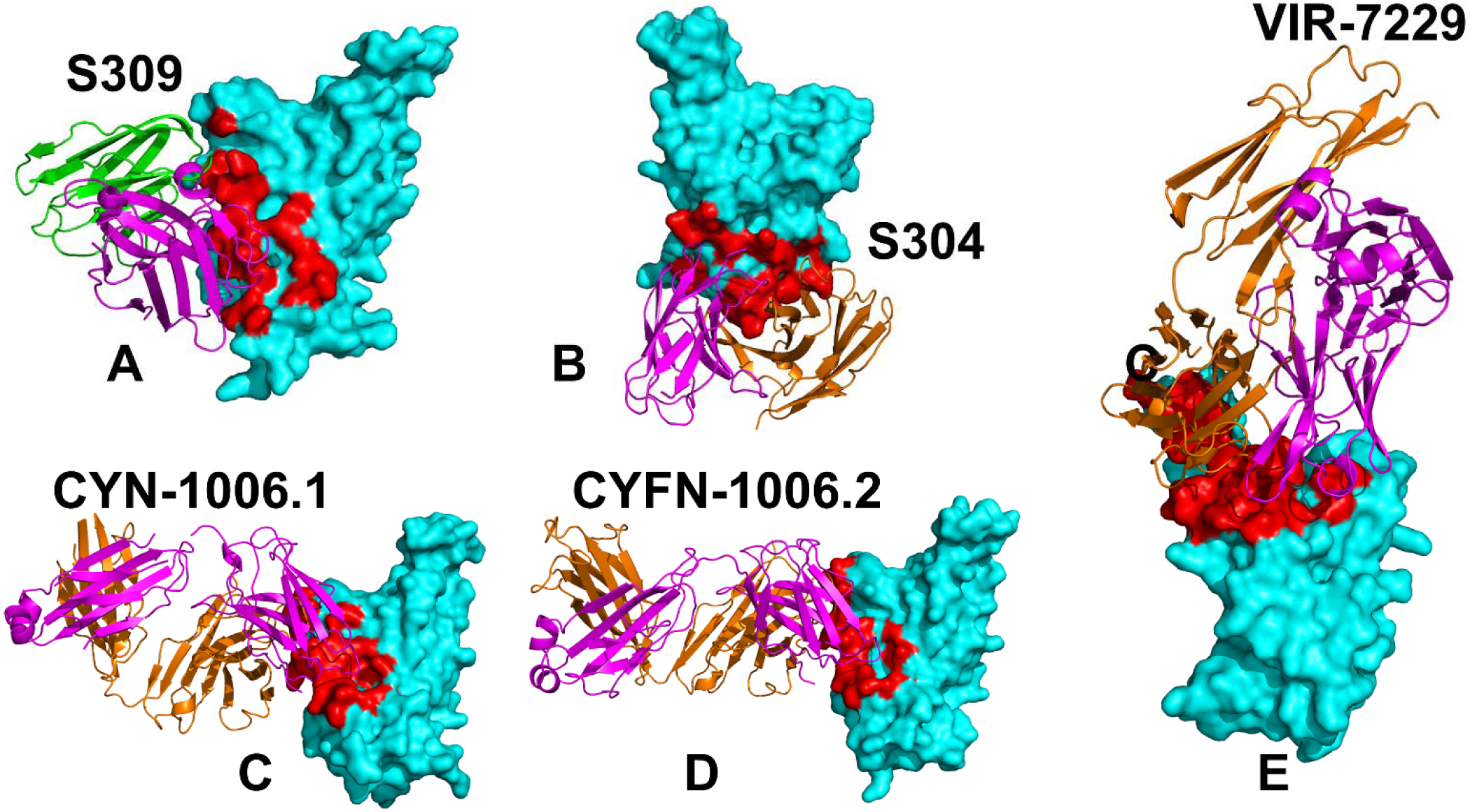
Structural organization of the SARS-CoV-2-RBD complexes with S309 (A), S304 (B), CYFN-1006.1 (C), CYFN-1006.2 (D) and VIR-7229 antibody (E). The S-RBD structure is shown in cyan surface. The heavy chains of antibodies are shown in orange ribbons and light chains are in magenta-colored ribbons. The binding epitope residues are shown in red surface.

S304 binds to a unique epitope located near residues F377, Y380, and T385 within the RBD (Figure 1B). This region is also outside the ACE2-binding motif but differs significantly from the S309 epitope [39]. S304 sterically interferes with the spike protein’s ability to adopt the "up" conformation necessary for ACE2 binding (Figure 1B) and achieves this through steric hindrance rather than direct competition or allosteric modulation [39]. The structures of the ternary Omicron RBD-hACE2-S304 complex, and the hACE2-bound Omicron spike with three reverse mutations (L371S, P373S, and F375S) by cryo-EM showed that S304 recognizes the inner face of the RBD buried in the down RBD configuration and its binding to the spike is conformation-dependent [40].

The two recently discovered antibodies CYFN1006-1 and CYFN1006-2 (Figure 1C,D) demonstrated consistent neutralization of all tested SARS-CoV-2 variants, outperforming SA55 [41]. These antibodies have binding epitopes overlapping with LY-CoV1404, REGN10987, and S309, located on the outer surface of the RBD (Figure 1C,D). A yeast-display system combined with a machine learning (ML)-guided approach for library design enabled an investigation of a larger number of antibody variants and the identification of a class 1 human antibody designated as VIR-7229, which targets the receptor-binding motif (RBM) potently neutralizing SARS-CoV-2 variants, including EG.5, BA.2.86, and JN.1 [42]. The structures of VIR-7229-bound to XBB.1.5 and EG.5 structures showed that the VIR-7229 interactions can accommodate both F456 and L456 in the corresponding genetic backgrounds and tolerate an extraordinary epitope variability exhibiting high barrier for the emergence of resistance, partly attributed to its high binding affinity [43]. Taken together, these findings underscore the complexity of the immune response to SARS-CoV-2 and highlight the importance of considering both epitope specificity and mutational escape profiles when evaluating antibody therapeutics or vaccine-induced immunity. CYFN1006-1 binds to the outer side of RBD with five of six complementary determining regions of CYFN1006-1 are engaged in RBD binding, involving a total of 17 RBD residues (Figure 1C,D) [41]. VIR-7229 binds to a structurally complex epitope within the RBM of the RBD, comprising 25 amino acid residues distributed across multiple regions of the RBD. These residues include 403, 405, 409 located near the N-terminal edge of the RBM [42]. Structural studies have revealed that VIR-7229 binding induces significant conformational changes in the RBD, particularly in residues 473–489 [42].

Computer simulations have emerged as indispensable tools for elucidating the molecular mechanisms of the S protein, its interactions with the ACE2 and its ability to evade neutralizing antibodies at the atomic level [44-48]. These computational approaches provide unparalleled insights into the structural and energetic factors that govern viral-host interactions and immune escape strategies. By employing advanced techniques such as molecular dynamics (MD) simulations and Markov state models (MSM), we have systematically mapped the conformational landscapes of Omicron subvariants like XBB.1 and XBB.1.5, as well as their complexes with ACE2 and antibodies [49]. Such studies have revealed intricate details about how these variants achieve enhanced receptor binding while simultaneously evading antibody-mediated immunity. Mutational scanning and binding analyses of XBB variants have identified key residues—including Y501, R498, Q493, L455F, and F456L—that contribute to epistatic interactions, strengthening ACE2 binding while conferring resistance to neutralizing antibodies [50,51]. Notably, convergent mutations such as F456L and F486P underscore the virus’s remarkable capacity to balance receptor affinity with immune evasion. Additionally, integrating AlphaFold2-based predictions with ensemble analyses of S protein-ACE2 complexes has enabled researchers to identify binding energy hotspots and epistatic networks involving critical residues like L455, F456, and Q493 in variants such as JN.1, KP.1, KP.2, and KP.3 [52]. These findings illustrate how the virus maintains high-affinity interactions with ACE2 while acquiring mutations that facilitate immune escape. Using MD simulations, ensemble-based deep mutational scanning of SARS-CoV-2 spike residues and binding free energy computations we examined mechanisms of broadly neutralizing antibodies : E1 group (BD55-3152, BD55-3546 and BD5-5840) and F3 group (BD55-3372, BD55-4637 and BD55-5514) [53]. Our analysis revealed the emergence of a small number of immune escape positions for E1 group antibodies that correspond to R346 and K444 positions in which the strong van der Waals and interactions act synchronously leading to the large binding contribution. A comprehensive structural and energetic analysis of the RBD complexes with neutralizing antibodies from four distinct groups (A-D), including group A LY-CoV016, group B AZD8895 and REGN10933, group C LY-CoV555, and group D antibodies AZD1061, REGN10987, and LY-CoV1404 identified key binding hotspots, and explored the evolutionary strategies employed by the virus to evade neutralization [54]. Previous studies also demonstrated that the S protein functions as an allosteric regulatory machine, leveraging its intrinsic flexibility to modulate receptor binding and immune evasion [55-58]. These studies reveal that antibody-escaping mutations often target structurally adaptable energy hotspots and allosteric effector centers, which are critical for maintaining the functional integrity of the S protein. Electrostatic interactions have emerged as a critical thermodynamic force governing the binding of the S protein to ACE2 and its resistance to antibodies [59-61]. The accumulation of positively charged residues on the RBD of many Omicron variants reflects evolutionary adaptations that enhance ACE2 binding while promoting immune evasion.

The evolution of SARS-CoV-2, particularly within the Omicron lineage, has been characterized by the emergence of subvariants with enhanced immune evasion and transmissibility, most recently JN.1, KP.2 and KP.3 variants [62]. JN.1 has evolved into multiple subvariants, each with distinct mutations that contribute to immune evasion and transmissibility (Supporting Information, Table S1). KP.2 that carries mutations R346T, F456L, and V1104L and KP.3 that features mutations R346T, L455S, F456L, Q493E, and V1104L share the F456L mutation and KP.3 has emerged as the most immune-evasive and fastest-growing JN.1 sublineage, largely due to the F456L mutation, which plays a critical role in the antibody escape [63]. F456L mutation enhances the binding potential of Q493E, leading to stronger receptor interactions and providing an evolutionary advantage for incorporating additional immune-evasive mutation [64,65]. Many studies suggested functionally balanced substitutions that optimize tradeoffs between immune evasion, high ACE2 affinity and sufficient conformational adaptability might be a common strategy of the virus evolution and serve as a primary driving force behind the emergence of new Omicron subvariants [66-68].

Experimental and computational studies collectively demonstrated that cross-neutralization against Omicron variants is governed by a complex and delicate interplay of multiple energetic factors and interaction contributions all of which shape the evolution of escape hotspots linked to antigenic drift and convergent evolution. The dynamic and energetic factors are closely tied to the evolution of immune escape hotspots that arise as a result of antigenic drift and convergent evolution. Such evolutionary adaptations allow the virus to evade immune pressure while maintaining its ability to bind effectively to the ACE2 receptor on host cells, thereby preserving its infectivity. The molecular mechanisms underlying these adaptations involve a fine-tuned balance between competing demands: enhancing immune evasion while avoiding significant losses in receptor-binding affinity. Moreover, the evolutionary trade-offs shaping the virus’s ability to balance immune evasion and ACE2 binding are highly nuanced and context-dependent. These trade-offs are influenced not only by the specific mutations present in the viral genome but also by the characteristics of the antibodies themselves. Different antibodies target distinct epitopes on the RBD, and their neutralization efficacy can vary widely depending on the precise mutational landscape of the variant in question. This antibody-dependent variability adds another layer of complexity to understanding how SARS-CoV-2 continues to adapt under selective pressures imposed by both natural immunity and vaccination. Cross-neutralization activity of antibodies against Omicron variants reflects a dynamic equilibrium shaped by multiple factors, including the energetic contributions of specific molecular interactions, the distribution of escape hotspots across the spike protein, and the selective pressures exerted by diverse antibody repertoires.

In this study, we expand upon existing research by delving into the molecular mechanisms of antibody binding through a comparative analysis of dynamic and energetic properties of S309, S304, GYFN-1006, and VIR-7229 antibody complexes with the RBD of the SARS-CoV-2 spike protein. Our approach integrates conformational dynamics, analysis of collective motions using Principal Component Analysis (PCA), along with systematic mutational profiling of RBD residues and rigorous binding free energy computations and residue-based energetic analysis of the RBD-antibody complexes. This analysis provides a detailed understanding of distinct molecular mechanisms and interactions that govern antibody recognition and neutralization.

Using MD simulations, we examined dynamic ensembles that represent the structural variability and flexibility of these systems and allowed us to examine how mutations in the RBD influence the stability and adaptability of the binding interfaces. By analyzing the trajectories, we identified key regions of the RBD that exhibit significant conformational changes upon antibody binding. These changes are critical for understanding how antibodies modulate the spike protein’s ability to interact with the ACE2 receptor while evading immune detection. Functional motions were further dissected using PCA, which revealed collective modes of motion that dominate the dynamics of the complexes. We show that residues located at the hinge regions of the RBD play a pivotal role in mediating transitions between open and closed conformations, thereby influencing both ACE2 binding affinity and antibody neutralization efficiency.

To systematically assess the impact of mutations on antibody binding, we conducted mutational scanning of RBD residues across the S-antibody complexes and generated mutational sensitivity heatmaps, which highlighted "escape hotspot" centers—regions where mutations significantly reduced antibody binding affinity. These hotspots often coincided with epitope residues directly contacted by the antibodies, but intriguingly, some non-contact residues also emerged as critical contributors due to their roles in maintaining the structural integrity of the binding interface. The identification of escape hotspots provides valuable insights into the evolutionary strategies employed by SARS-CoV-2 variants to evade neutralizing antibodies. To rigorously quantify the binding affinities of the S-antibody complexes, we employed the Molecular Mechanics/Generalized Born Surface Area (MM-GBSA) approach. This method combines molecular mechanics calculations with implicit solvation models to estimate the free energy contributions of individual residues to the overall binding affinity. Through residue-based energy decomposition, we identified key residues that contribute disproportionately to the stability of the binding interfaces. Mutational profiling and binding affinity analysis identifies key residues crucial for antibody binding underscoring their roles as evolutionary "weak spots" that balance viral fitness and immune evasion.

The results of this study dissect distinct energetic mechanisms of binding and importance of targeting conserved residues and diverse epitopes to counteract viral resistance. In particular, we show that broad-spectrum antibodies CYFN1006 and VIR-7229, which maintain efficacy across multiple variants, can achieve neutralization by targeting convergent evolution hotspots R346 and F456 while enabling tolerance to mutations in these positions through structural adaptability and compensatory interactions at the binding interface. The results of this study underscore the diversity of binding mechanisms employed by different antibodies and molecular basis for high affinity and excellent neutralization activity of the latest generation of antibodies.

## Materials and Methods

### Molecular Dynamics Simulations

The crystal and cryo-EM structures of the Omicron RBD-complexes are obtained from the Protein Data Bank [69]. For simulated structures, hydrogen atoms and missing residues were initially added and assigned according to the WHATIF program web interface [70]. The missing regions are reconstructed and optimized using template-based loop prediction approach ArchPRED [71]. The side chain rotamers were refined and optimized by SCWRL4 tool [72]. The protonation states for all the titratable residues of the antibody and RBD proteins were predicted at pH 7.0 using Propka 3.1 software and web server [73,74]. The protein structures were then optimized using atomic-level energy minimization with composite physics and knowledge-based force fields implemented in the 3Drefine method [75,76]. NAMD 2.13-multicore-CUDA package [77] with CHARMM36 force field [78] was employed to perform 1µs all-atom MD simulations for the RBD-antibody complexes. The structures of the complexes were prepared in Visual Molecular Dynamics (VMD 1.9.3) [79] and with the CHARMM-GUI web server [80,81] using the Solutions Builder tool. Hydrogen atoms were modeled onto the structures prior to solvation with TIP3P water molecules [82] in a periodic box that extended 10 Å beyond any protein atom in the system. To neutralize the biological system before the simulation, Na+ and Cl− ions were added in physiological concentrations to achieve charge neutrality, and a salt concentration of 150 mM of NaCl was used to mimic physiological concentration. All Na+ and Cl− ions were placed at least 8 Å away from any protein atoms and from each other. MD simulations are typically performed in an aqueous environment in which the number of ions remains fixed for the duration of the simulation, with a minimally neutralizing ion environment or salt pairs to match the macroscopic salt concentration [83]. All systems were subjected to a minimization protocol consisting of two stages. First, minimization was performed for 100,000 steps with all the hydrogen-containing bonds constrained and the protein atoms fixed. In the second stage, minimization was performed for 50,000 steps with all the protein backbone atoms fixed and for an additional 10,000 steps with no fixed atoms. After minimization, the protein systems were equilibrated in steps by gradually increasing the system temperature in steps of 20 K, increasing from 10 K to 310 K, and at each step, a 1ns equilibration was performed, maintaining a restraint of 10 kcal mol−1 Å−2 on the protein Cα atoms. After the restraints on the protein atoms were removed, the system was equilibrated for an additional 10 ns. Long-range, non-bonded van der Waals interactions were computed using an atom-based cutoff of 12 Å, with the switching function beginning at 10 Å and reaching zero at 14 Å. The SHAKE method was used to constrain all the bonds associated with hydrogen atoms. The simulations were run using a leap-frog integrator with a 2 fs integration time step. The ShakeH algorithm in NAMD was applied for the water molecule constraints. The long-range electrostatic interactions were calculated using the particle mesh Ewald method [84] with a cut-off of 1.0 nm and a fourth-order (cubic) interpolation. The simulations were performed under an NPT ensemble with a Langevin thermostat and a Nosé–Hoover Langevin piston at 310 K and 1 atm. The damping coefficient (gamma) of the Langevin thermostat was 1/ps. In NAMD, the Nosé–Hoover Langevin piston method is a combination of the Nosé–Hoover constant pressure method [85] and piston fluctuation control implemented using Langevin dynamics [86,87]. An NPT production simulation was run on equilibrated structures for 1µs keeping the temperature at 310 K and a constant pressure (1 atm).

### Mutational Scanning of the Binding Interfaces for the the SARS-CoV-2 S Protein Complexes with Antibodies

To understand the molecular mechanisms underlying the interactions between the SARS-CoV-2 S-RBD and neutralizing Abs, we conducted a comprehensive mutational scanning analysis of the binding epitope residues. This approach systematically evaluated the effects of mutations on protein stability and binding free energy, providing insights into the structural and energetic determinants of RBD-antibody interactions. Each binding epitope residue in the RBD-antibody complexes was systematically mutated using all possible amino acid substitutions. The corresponding changes in protein stability and binding free energy were computed using the BeAtMuSiC approach [88-90]. This method relies on statistical potential that describe pairwise inter-residue distances, backbone torsion angles, and solvent accessibility. The BeAtMuSiC approach evaluates the impact of mutations on both the strength of interactions at the protein-protein interface and the overall stability of the complex. The binding free energy of a protein-protein complex is expressed as the difference between the folding free energy of the complex and the folding free energies of the individual binding partners:

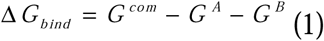

The change of the binding energy due to a mutation was calculated then as

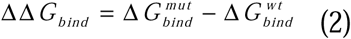

To ensure robust results, we leveraged rapid calculations based on statistical potentials, computing ensemble-averaged binding free energy changes using equilibrium samples from molecular dynamics (MD) simulation trajectories. The binding free energy changes were averaged over 10,000 equilibrium samples for each system studied. We used 1000 ns of equilibrated trajectory data for each system, with snapshots collected at 100 ps intervals.

### Binding Free Energy Computations of the SARS-CoV-2 S Protein Complexes with Antibodies

We employed the Molecular Mechanics/Generalized Born Surface Area (MM/GBSA) method [91,92] for binding free energy calculations of the S-antibody complexes To ensure robust sampling, equilibrium trajectories were extracted from the production phase of the MD simulations. Specifically, we used 1000 ns of equilibrated trajectory data for each system, with snapshots collected at 100 ps intervals. This approach provided a total of 10,000 frames per system for binding free energy calculations. Additionally, we conducted an energy decomposition analysis to evaluate the contribution of each amino acid during the binding of RBD to Antibodies [93,94].

The binding free energy for the RBD-Antibody complex was obtained using:

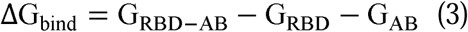

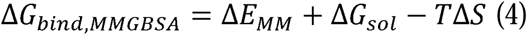

where ΔE_MM_ is total gas phase energy (sum of ΔEinternal, ΔEelectrostatic, and ΔEvdw); ΔGsol is sum of polar (ΔGGB) and non-polar (ΔGSA) contributions to solvation. Here, G _RBD–ANTIBODY_ represent the average over the snapshots of a single trajectory of the complex, G_RBD_ and G_ANTIBODY_ corresponds to the free energy of RBD and antibody respectively.

The polar and non-polar contributions to the solvation free energy is calculated using a Generalized Born solvent model and consideration of the solvent accessible surface area [95]. MM-GBSA is employed to predict the binding free energy and decompose the free energy contributions to the binding free energy of a protein–protein complex on per-residue basis. The binding free energy with MM-GBSA was computed by averaging the results of computations over 10,000 samples from the equilibrium ensembles. We used 1000 ns of equilibrated trajectory data for each system, with snapshots collected at 100 ps intervals. This approach provided a total of 10,000 frames per system for binding free energy calculations. Single trajectory protocol uses one trajectory of the RBD-antibody complex, reducing noise by canceling out intermolecular energy contributions. This protocol is suitable when significant structural changes upon binding are not expected. We employed a single-trajectory protocol due to its lower noise and applicability to systems with minimal structural reorganization upon binding. Entropy contributions are typically excluded from calculations because the entropic differences in relative binding affinities are expected to be small for minor mutational changes [96,97]. MM/GBSA calculations were performed using the MMPBSA.py module in AMBER, with the dielectric constant set to 1 for the solute and 80 for the solvent in the AmberTools21 package [98] and gmx_MMPBSA, a new tool to perform end-state free energy calculations from CHARMM and GROMACS trajectories [99].

## Results

### Structural Analysis of Binding Epitopes and MD Simulations of the SRBD-Complexes with Antibodies

All-atom MD simulations were conducted for the S-RBD complexes with a panel of studied antibodies to explore their conformational landscapes and identify specific dynamic signatures induced by antibody binding. The primary objective of this study was to investigate the dynamic and energetic contributions of RBD residues, as these residues play a pivotal role in mediating interactions with both host cell receptors and neutralizing antibodies. However, it is important to acknowledge that the flexibility of antibody residues—particularly within the complementarity-determining regions (CDRs)—could also influence the overall binding mechanism. While antibodies are generally more rigid compared to the RBD, subtle variations in their dynamics may contribute to the fine-tuning of antigen recognition. Nevertheless, given the central focus of this work on the RBD, the dynamic behavior of antibodies was not explored in depth.

The epitope recognized by S309 is located on the outer surface of the RBD, which becomes exposed when the spike protein adopts the "up" conformation required for receptor engagement (Figure 1). This accessibility ensures that S309 can effectively bind to the RBD during viral entry. The S309 epitope involves residues spanning three key regions. Residues 334–346 stretch includes highly conserved residues that are critical for stabilizing the interaction between S309 and the RBD. Residues 354–359 contribute to the hydrophobic and electrostatic interactions that enhance binding affinity. Structural loop (443–450) plays a role in mediating antibody recognition. Notably, residues L441 and K444 are central to the interaction, forming hydrogen bonds and van der Waals contacts with S309 (Figure 2A,B). By targeting a region near the ACE2-binding site, S309 effectively blocks viral entry while maintaining broad-spectrum neutralization activity across multiple variants. In contrast, the S304 antibody recognizes a distinct epitope on the RBD that involves residues from multiple structural elements, including α-helices, β-strands, and flexible loops. This epitope is structurally complex and contributes to the formation of the conserved RBD β-sheet. The S304 epitope comprises residues 369–392, which are part of two α-helices and an intervening β-strand (Figure 2C,D). These regions participate in the formation of the structurally conserved RBD β-sheet, underscoring the stability of the interaction. The binding epitope of S304 includes the following key residues Y369, N370, F374, S375, F377, K378, 379, 380–386, 390, 392 that form a dense network of interactions, including hydrogen bonds, van der Waals forces, and hydrophobic packing.

**Figure 2.**
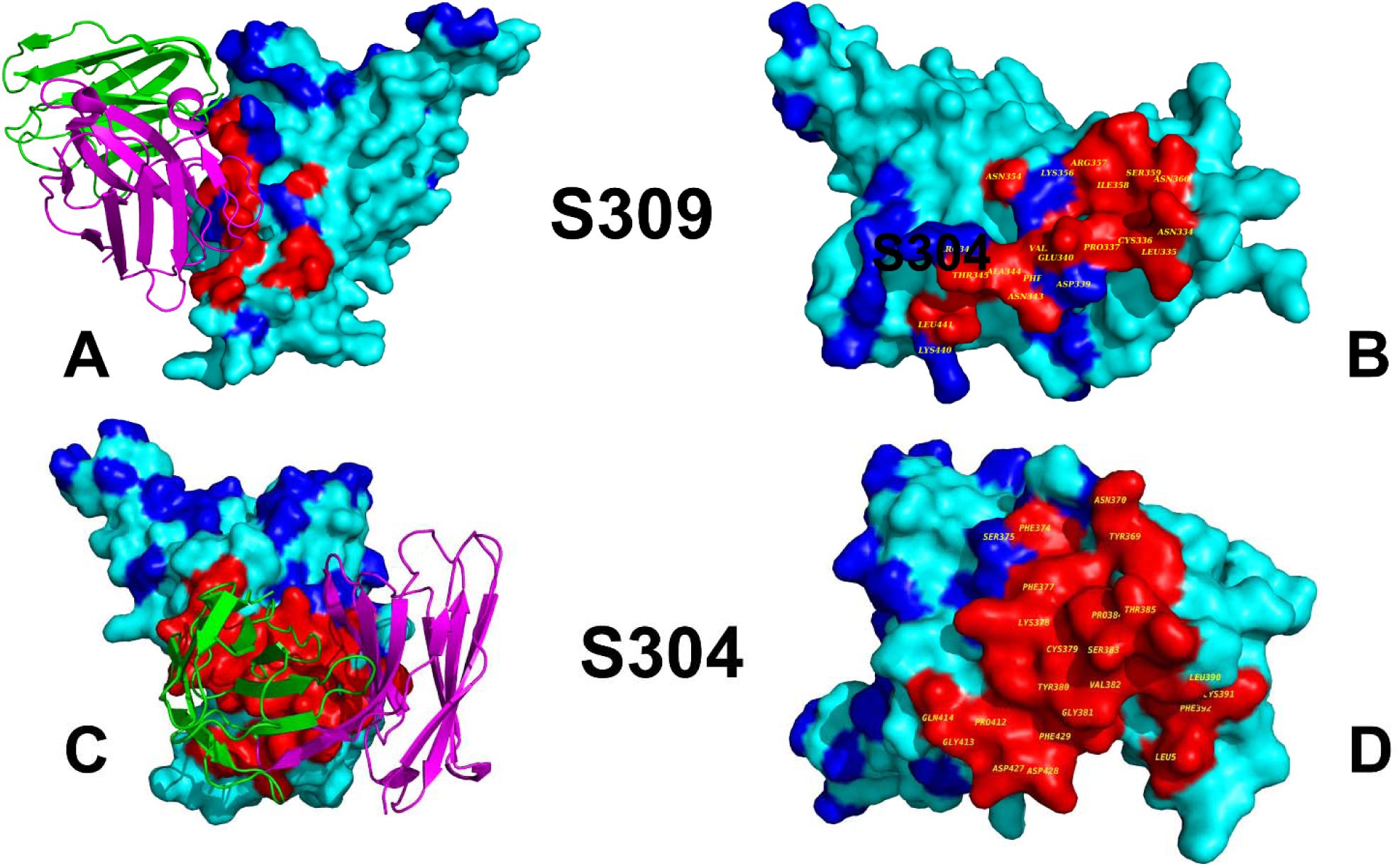
Structural organization of the RBD complexes and binding epitopes of S304 (A,B) and S304 antibody (C,D). The S-RBD structure is shown in cyan surface. The heavy chains of antibodies are in orange ribbons and the light chains are in magenta ribbons. (B) The RBD surface and binding epitope for S309-RBD complex. RBD is shown in cyan surface. The binding epitope residues are in red surface (N334, L335, P337, D339, E340, N343, A344, T345, R346, F347, N354 K356, R357, S359, L441, and K444). (D) The RBD surface and binding epitope for S304-RBD complex. RBD is shown in cyan surface. The binding epitope residues are in red surface (Y369, N370, F374, F377, K378, C379, Y380, G381, V382, S383, P384, T385, K386, L390, F392, P412, G413, D427, T430, L517) The key sites of Omicron XBB, BA.2.86, and JN.1 lineages are shown in blue surface (residues 339, 346, 356, 371, 373, 375 376, 403, 405, 408, 417, 440, 444, 445, 446, 450, 452, 455, 456, 460, 475, 477, 478, 481, 484, 486, 493, 498, 501, 505).

Additionally, loop residues 411–414 and 427–430 contribute to the epitope, providing flexibility and adaptability during antibody binding (Figure 2C,D). Residue 517, which is embedded in the hydrophobic core of the epitope, plays a particularly important role in stabilizing interfacial interaction. The involvement of both structured elements (α-helices and β-strands) and flexible loops highlights the cooperative nature of S304 binding. This dual engagement ensures strong and specific interactions, even in the presence of minor mutations in the RBD. While both S309 and S304 are potent neutralizing antibodies, their epitopes differ significantly in terms of location, composition, and functional implications. S309 binds near the ACE2-binding site, allowing it to block receptor engagement through steric hindrance. S304 targets a more peripheral region of the RBD, involving residues that contribute to the structural integrity of the RBD β-sheet. While the S309 epitope involves conserved residues 334-346 and a structural loop (443–450) near the ACE2 interface, the S304 epitope encompasses a broader range of structural elements, including α-helices, β-strands, and flexible loops. The S309 epitope is highly conserved, making it less susceptible to mutations (Figure 2A,B). While the S304 epitope also includes conserved residues, its reliance on flexible loops may render it more sensitive to mutational changes (Figure 2C,D).

We performed a detailed examination of the conformational ensembles of the RBD protein. This analysis elucidates the effects of RBD mutations on its dynamics and molecular determinants of RBD-antibody binding—a key driver of viral evolution. To characterize the dynamic flexibility of the RBD when bound to different antibody groups, we performed root-mean-square fluctuation (RMSF) analysis on equilibrated MD trajectories (Figure 3). First, we compared the RMSF profiles of the RBD-S309 complexes obtained from MD simulations of four different RBD-S309 structures for different Omicron variants BA.1, XBB.1, BQ.1.1 and BN.1 (Figure 3A). The use of multiple systems allowed for a comparative analysis of RBD dynamics across variants, highlighting both shared and unique features. The RMSF profiles of the RBD-S309 complexes revealed a high degree of similarity across the four Omicron subvariants, reflecting the conserved structural core of the RBD. However, notable differences were observed in specific regions. The central β-sheet and α-helices of the RBD exhibited low RMSF values, indicating minimal flexibility. These regions are critical for maintaining the overall structural integrity of the RBD. Two key regions displayed higher flexibility. Residues 355–375 showed moderate fluctuations, likely due to its proximity to the S309 epitope and involvement in mediating interactions with the antibody. 470–490 Loop exhibited the highest RMSF values, underscoring its role as a highly dynamic element within the RBD. The 470–490 loop is a well-characterized conformationally dynamic region of the RBD. The loop is located near the ACE2-binding interface and often contributes to stabilizing the RBD-ACE2 interaction. The loop is also implicated in mediating interactions with neutralizing antibodies, including S309. In the RMSF profiles, the 470–490 loop consistently showed elevated flexibility across all four variants, although subtle differences were observed. XBB.1 and BQ.1.1 variants displayed higher flexibility, suggesting that mutations in these subvariants may enhance the loop’s adaptability to accommodate antibody binding. The increased flexibility of the 470–490 loop allows it to adjust its conformation to optimize interactions with S309. This adaptability may enhance the stability of the RBD-S309 complex while potentially interfering with ACE2 binding.

**Figure 3.**
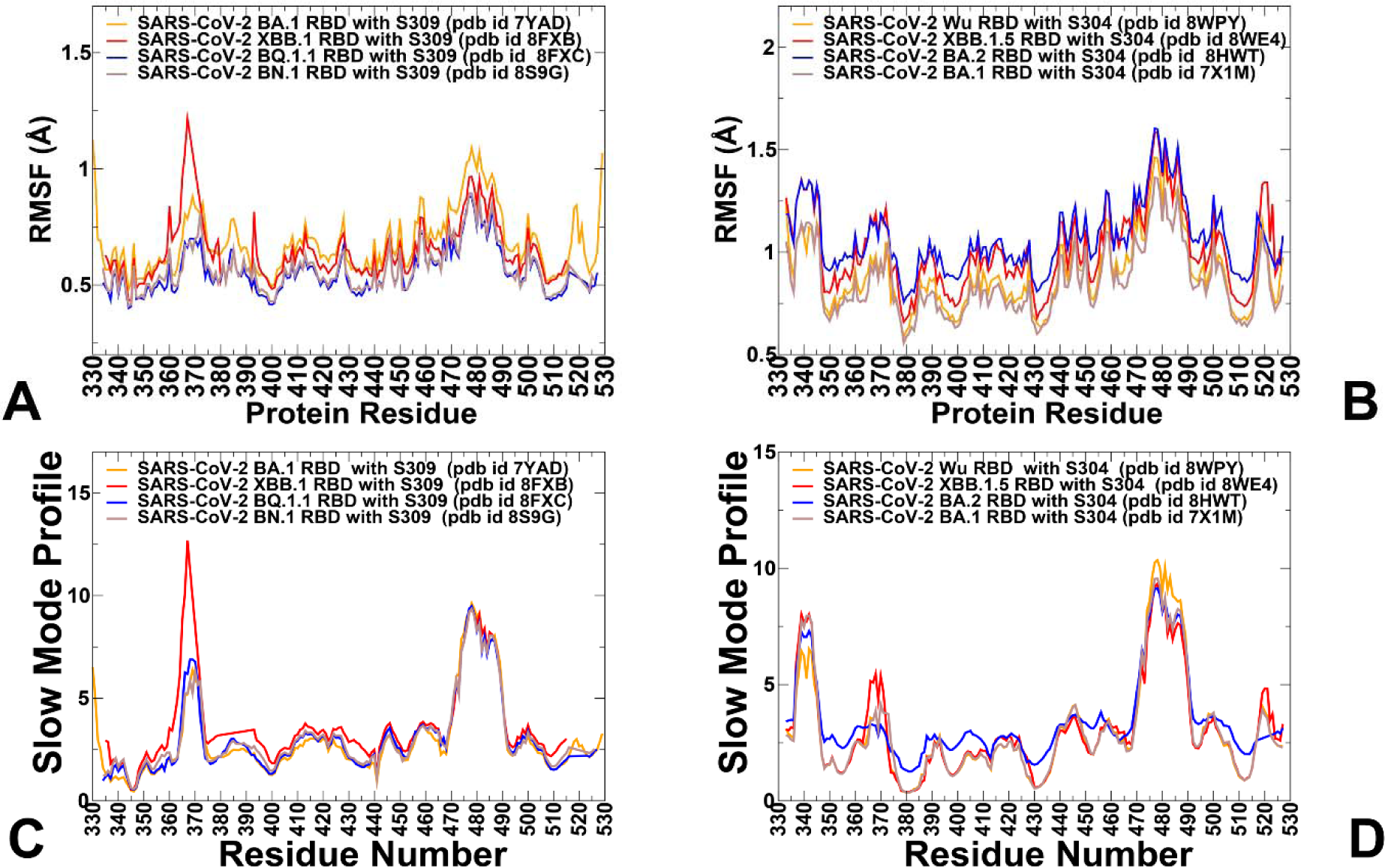
Conformational dynamics profiles obtained from simulations of the RBD-antibody complexes. (A) The RMSF profiles for the RBD residues obtained from MD simulations of the S-RBD complexes with S309 using different structures : BA.1 RBD, pdb id 7YAD (in orange lines), XBB.1 RBD, pdb id 8FXB (in red lines), BQ.1.1 RBD, pdb id 8FXC (in blue lines) and BN1RBD pdb id 8S9G (in light brown lines). (B) The RMSF profiles for the RBD residues obtained from MD simulations of the S-RBD complexes with S304 antibody using different structures : Wu-WT, pdb id 8WPY (in orange lines), XBB.1.5 RBD, pdb id 8WE4 (in red lines), BA2 RBD, pdb id 8HWT (in blue lines) and BA.1 RBD, pdb id 7X1M (in light brown lines). (C) The slow mode profile averaged over 10 lowest frequency modes obtained from MD simulations of RBD-S309 complexes using different structures : BA.1 RBD, pdb id 7YAD (in orange lines), XBB.1 RBD, pdb id 8FXB (in red lines), BQ.1.1 RBD, pdb id 8FXC (in blue lines) and BN1RBD pdb id 8S9G (in light brown lines). (D) The slow mode profile averaged over 10 lowest frequency modes obtained from MD simulations of RBD-S304 complexes using different structures : Wu-WT, pdb id 8WPY (in orange lines), XBB.1.5 RBD, pdb id 8WE4 (in red lines), BA2 RBD, pdb id 8HWT (in blue lines) and BA.1 RBD, pdb id 7X1M (in light brown lines).

S304 antibody interacts with specific regions of the RBD, including residues 369–392 (two α-helices and an intervening β-strand), residues 515–517 (part of the structurally conserved RBD β-sheet), and loop residues 411–414 and 427–430. The RMSF values for residues 369–392 are relatively low, indicating minimal flexibility (Figure 3B) These residues form the core of the epitope and play a pivotal role in stabilizing the interaction between S304 and the RBD. Residues Y369, N370, F374, S375, F377, K378, 379, 380–386, 390, and 392 form a dense network of interactions, including hydrogen bonds, van der Waals forces, and hydrophobic packing. These interactions further reduce flexibility and enhance binding affinity. The RMSF values for loop residues 411–414 and 427–430 are higher compared to the structured core, indicating significant flexibility. This flexibility allows these loops to adjust their conformation to optimize interactions with the antibody. The loops play a crucial role in mediating antibody recognition by forming hydrogen bonds and van der Waals contacts with S304. Their adaptability ensures that the interaction remains stable even in the presence of minor mutational changes. S309 binding stabilizes the core of the RBD, locking it into a rigid conformation that is less conducive to large-scale motions. This stabilization indirectly restricts the flexibility of the 470–490 loop S304 engages a broader and more diverse set of residues, requiring the RBD to adopt a more dynamic conformation. This leads to increased flexibility in the 470–490 loop to accommodate the antibody’s binding requirements. The increased flexibility of the 470–490 loop in the S304-bound state highlights its role in adaptive binding (Figure 3A,B). This adaptability ensures that S304 can effectively neutralize diverse variants despite antigenic drift. Conversely, the reduced flexibility of the loop in the S309-bound state reflects the antibody’s ability to stabilize the RBD in a conformation that sterically hinders ACE2 binding, thereby preventing viral entry.

We also characterized essential motions and determined the hinge regions in the RBD-S309 and RBD-S304 complexes (Figure 3C,D) using principal component analysis (PCA) of trajectories using the CARMA package [100] The local minima along these profiles are typically aligned with the immobilized in global motions hinge centers, while the maxima correspond to the moving regions undergoing concerted movements leading to global changes in structure. The low-frequency ‘soft modes’ are characterized by their cooperativity and there is a strong relationship between conformational changes and the ‘soft’ modes of motions intrinsically accessible to protein structures [101,102]. The slow mode profile computed by averaging essential motions over the ten lowest modes revealed key hinge positions corresponding to T346, F375, L441 and D467 residues (Figure 3C). Group E antibodies are particularly sensitive to mutations at residues G339, T345, and R346, which are located near the N-terminal region of the RBD. These residues play a critical role in stabilizing the RBD structure and facilitating interactions with certain antibodies. L441 is central to the interaction, forming hydrogen bonds and van der Waals contacts with S309. The regions that undergo large movements in slow modes are regions 362-375 and 470-490 (Figure 3C). Interestingly, a large fraction of the RBD becomes immobilized in slow motions upon S309 binding. This rigidity results from the antibody’s ability to induce a more constrained conformation that is less conducive to large-scale motions. The stabilization of the RBD core and reduction in flexibility highlight the robust nature of S309 binding and its effectiveness in neutralizing the virus.

In contrast to the RBD-S309 complex, the slow mode profile of the RBD-S304 complex features a more dynamic behavior, characterized by hinge regions that are aligned with positions 374–385 and 429–436, indicating a broader distribution of pivot points compared to the RBD-S309 complex (Figure 3D). Several regions exhibit enhanced mobility in the presence of S304, including residues 360–380, 430-450 and 468–494 The RBD-S304 complex exhibits more fluctuating behavior in global modes, as evidenced by a more distributed pattern of motion across multiple regions (Figure 3D). The enhanced flexibility of regions such as 360–380 and 430–450 suggests cooperative motions that enable the RBD to adopt multiple conformations during S304 binding. The slow mode profiles reveal distinct characteristics between the S309 and S304 complexes. The RBD-S309 complex exhibits greater rigidity, with fewer regions undergoing large-scale motions. In contrast, the RBD-S304 complex displays increased flexibility, particularly in regions near the epitope. The hinge sites in the RBD-S309 complex are more localized, whereas those in the RBD-S304 complex are more broadly distributed, reflecting the different binding mechanisms of the two antibodies. The increased flexibility observed in the RBD-S304 complex allows the RBD to adapt to mutations and maintain interactions with the antibody. However, this adaptability may also make it more susceptible to escape mutations in flexible regions.

The results on conformational dynamics reveal key insights into the flexibility and structural behavior of the SARS-CoV-2 RBD when interacting with S309 and S304. The central β-sheet and α-helices of the RBD exhibit low RMSF values, indicating minimal flexibility and highlighting their role in maintaining structural integrity. Two regions show higher flexibility: residues 355–375 and the 470–490 loop. The latter is particularly dynamic, located near the ACE2-binding interface, and plays a critical role in stabilizing RBD-ACE2 interactions and mediating antibody binding. S309 binding stabilizes the core of the RBD, reducing its flexibility and restricting large-scale motions. This stabilization indirectly limits the flexibility of the 470–490 loop, sterically hindering ACE2 binding and preventing viral entry. S304 binding requires the RBD to adopt a more dynamic conformation, increasing the flexibility of the 470–490 loop. This adaptability allows S304 to effectively neutralize diverse variants despite antigenic drift.

CYFN-1006 binds to a distinct epitope on the RBD that includes residues 339–348, 354–356, 399, 440–446, 450, and 499–500 (Figure 1E). Conserved core residues N343, A344, T345 are critical for forming hydrogen bonds with the antibody. Y107, A108, S114 of CDRL3 and D112A, and W112 of CDRH3 can form 7 pairs of hydrogen bonds with RBD residues N343, A344, and T345 [41]. Residues 339–348 form part of the core epitope and contributes significantly to the binding interface. N354, K356 are engaged in interactions with V30 of CDRL1, while K440, L441, P445, P499 are embedded in the hydrophobic patch formed by CDRH1 and CDRH3, these residues stabilize the hydrophobic core of the interface. S446, K444, T346 form hydrogen bonds with D109-W112 of CDRH3 further strengthening the interaction [41]. The RMSF profiles for the RBD in complex with CYFN-1006 revealed that residues 339–348 forming part of the conserved core epitope and exhibit low RMSF values, indicating minimal flexibility. Residues 354–356 show moderate flexibility, likely due to their proximity to the antibody’s complementarity-determining regions (CDRs) and involvement in mediating interactions with CYFN-1006. Residues 440–446 exhibit low RMSF values, reflecting the effect of binding. The flexible loop region 470-490 continues to be the most dynamic region in the complex (Figure 4A).

**Figure 4.**
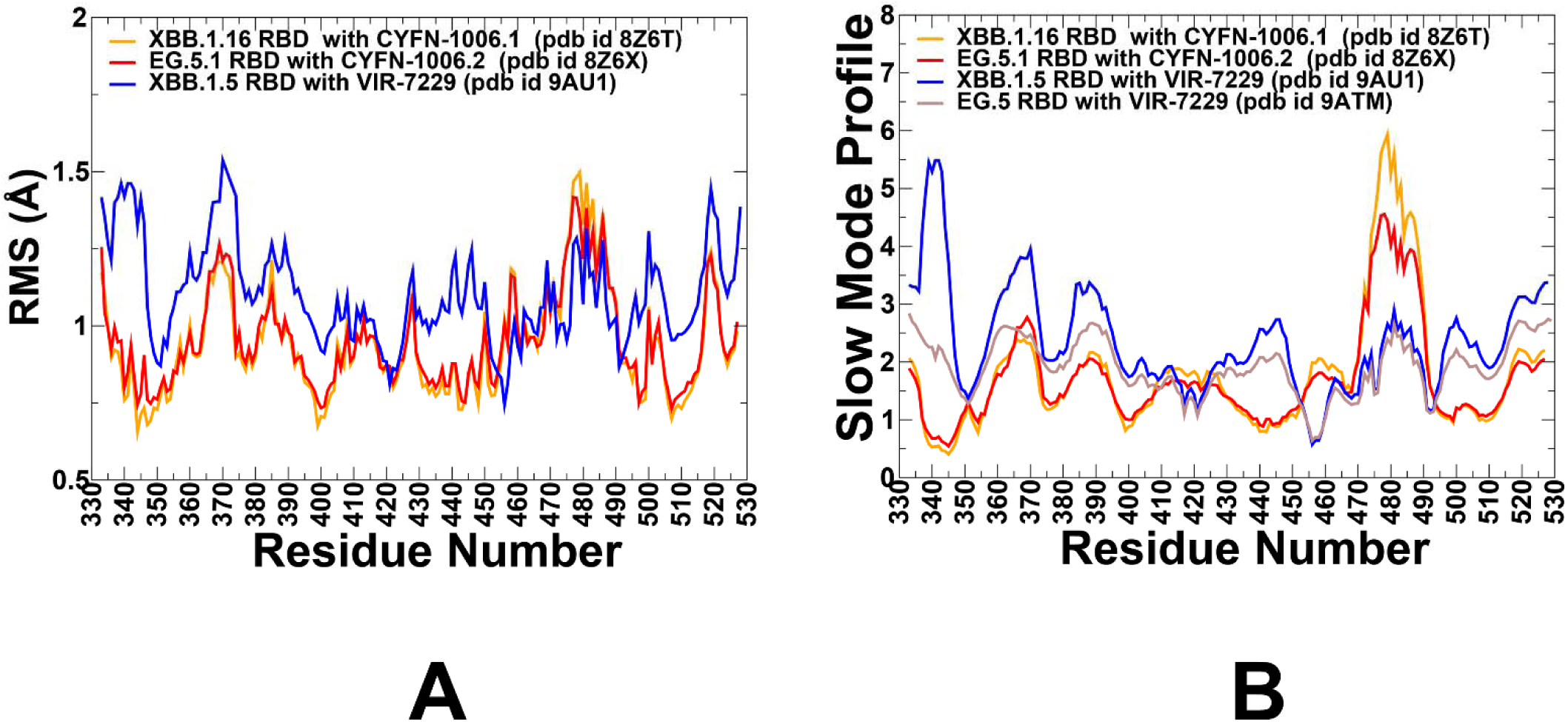
Conformational dynamics profiles obtained from simulations of the RBD-antibody complexes. (A) The RMSF profiles for the RBD residues obtained from MD simulations of the S-RBD complexes with CYFN-1006.1 (XBB.1.16 RBD, pdb id 8Z6T, in orange lines), CYFN-1006.2 (EG.5.1 RBD, pdb id 8Z6X, in red lines) and VIR-7229 (XBB.1.5 RBD, pdb 9AU1, in blue lines). (B) The slow mode profiles for the RBD residues obtained by averaging over 10 lowest frequency modes from MD simulations of the S-RBD complexes with CYFN-1006.1 (XBB.1.16 RBD, pdb id 8Z6T, in orange lines), CYFN-1006.2 (EG.5.1 RBD, pdb id 8Z6X, in red lines), VIR-7229 (XBB.1.5 RBD, pdb 9AU1, in blue lines) and VIR-7229 (EG.5 RBD, pdb id 9ATM, in brown lines).

VIR-7229 targets a broader and more structurally complex epitope on the RBD, including residues 403–417, 420–421, 453–460, 473–477, 487, 489, 493, and 505. Residues 415–417, 420–421 are central residues involved in direct interactions with VIR-7229. Residues 453–460 form a stretch of residues forming part of the hydrophobic core of the epitope. Residues 473–477, 487, 489, 493, 505 contribute to shape complementarity and polar interactions. 13 out of these 25 residues participate in binding to human ACE2, underscoring the overlap between the VIR-7229 epitope and the ACE2-binding interface [42]. The RMSF profiles for the RBD in complex with VIR-7229 revealed that residues 403–417 show moderate flexibility due to its involvement in direct interactions with VIR-7229 (Figure 4A). At the same time, region 425-460 exhibits elevated RMSF values, underscoring its role in mediating interactions with both ACE2 and antibodies. Loop Residues 473–477 exhibits moderately elevated RMSF values, underscoring its role in mediating interactions as VIR-7229 binding induces a significant rearrangement of these residues [42].

The analysis of slow mode profiles revealed the unique feature of functional motions for VIR-7229 complex (Figure 4B). In this case, the moving regions correspond to residues 355-375 while the rest of RBD residues experience only small movements including residues 470-490. A unique feature of VIR-7229 binding is its reliance on backbone-mediated interactions rather than side-chain contacts. This characteristic is reflected in the slow mode profiles of XBB.1.5 RBD complex as many hydrogen bonds formed between VIR-7229 and the RBD involve backbone atoms of residues such as N417, L455, R457, and K458 VIR-7229 forms four hydrogen bonds with the backbone of residues L455, R457, and K458 These positions and regions 417-422 and 455-460 become associated with the broad hinge points anchoring the RBD in the complex with VIR-7229 (Figure 4B). Hence, the slow mode analysis reveals that VIR-7229 induces significant rigidification of key regions within the RBD, particularly residues 455–460, which include the mutated position 456. Residues 455–460 become broad hinge points that anchor the RBD in the complex with VIR-7229. Structural studies also showed that VIR-7229 makes persistent contacts for both F456 (XBB.1.5) and L456 (EG.5) [42] and therefore can be effective against Omicron variants featuring mutations in these positions. We also observed that RBD regions 350-370 and 38-395 may undergo appreciable displacements in global motions, indicating that the ability of dynamic adjustments of the RBD to bind VIR-7229 (Figure 4B). Despite the F456L mutation in EG.5, the slow mode profile of the RBD-VIR-7229 complex remains largely unchanged. The region 455–460 continues to exhibit reduced flexibility, indicating that the antibody’s binding mode accommodates both F456 and L456 without compromising stability or altering location of the hinge points (Figure 4B). The ability of VIR-7229 to tolerate both F456 (found in XBB.1.5) and L456 (found in EG.5) can be attributed to its unique binding mechanism, which minimizes dependence on specific side-chain conformations and instead relies on backbone-mediated interactions. The antibody forms four hydrogen bonds with the backbone atoms of residues such as L455, R457, and K458, which stabilize the interaction regardless of whether F456 or L456 is present. This characteristic minimizes its dependence on the precise chemical properties of the side chains at position 456 (e.g., phenylalanine in F456 or leucine in L456). The slow mode analysis further supports this resilience by revealing how VIR-7229 immobilizes key regions of the RBD, including residues 455–460, locking F456 or L456 in the optimal binding position and preserving the character of functional motions. These hinge points anchor the RBD and stabilize the interaction, regardless of whether F456 or L456 is present. However, the rigidification induced by VIR-7229 binding affects different regions of the RBD depending on whether F456 or L456 is present. For F456, the rigidification is more localized around the core epitope (e.g., residues 455–460), while peripheral regions such as 350–370 and 380–395 retain some flexibility. These regions may undergo cooperative motions to facilitate binding. For L456, the reduced bulkiness of leucine leads to a broader suppression of functional displacements, as the smaller residue cannot stabilize the same degree of structural flexibility. The reduced rigidity of the hinge region suppresses these motions, resulting in a more static conformation of the EG.5 RBD in the complex. These findings highlight the adaptability of VIR-7229 to mutations of convergent evolution hotspots while also underscoring the nuanced impact of such mutations on RBD dynamics.

### Mutational Profiling of Protein Binding Interfaces

Using the conformational ensembles of the RBD complexes, we embarked on structure-based mutational analysis of the S protein binding with Abs. To provide a systematic comparison, we constructed mutational heatmaps for the RBD interface residues of the S complexes with S309, S304, CYFN1006-1, CYFN1006-2 and VIR-7229 antibodies. The mutational scanning of receptor-binding domain (RBD) residues in complexes with the S309 antibody provides critical insights into the structural and functional determinants of their interaction. Mutational scanning identified the following residues as critical for binding S309 antibody: N334, L335, P337, D339, T345, R346, and L441 (Figure 5, Supporting Information, Datasets S1-S4). N334 is part of the helical region (residues 337–344) that interacts with the complementarity-determining region (CDR) loops of S309.

**Figure 5.**
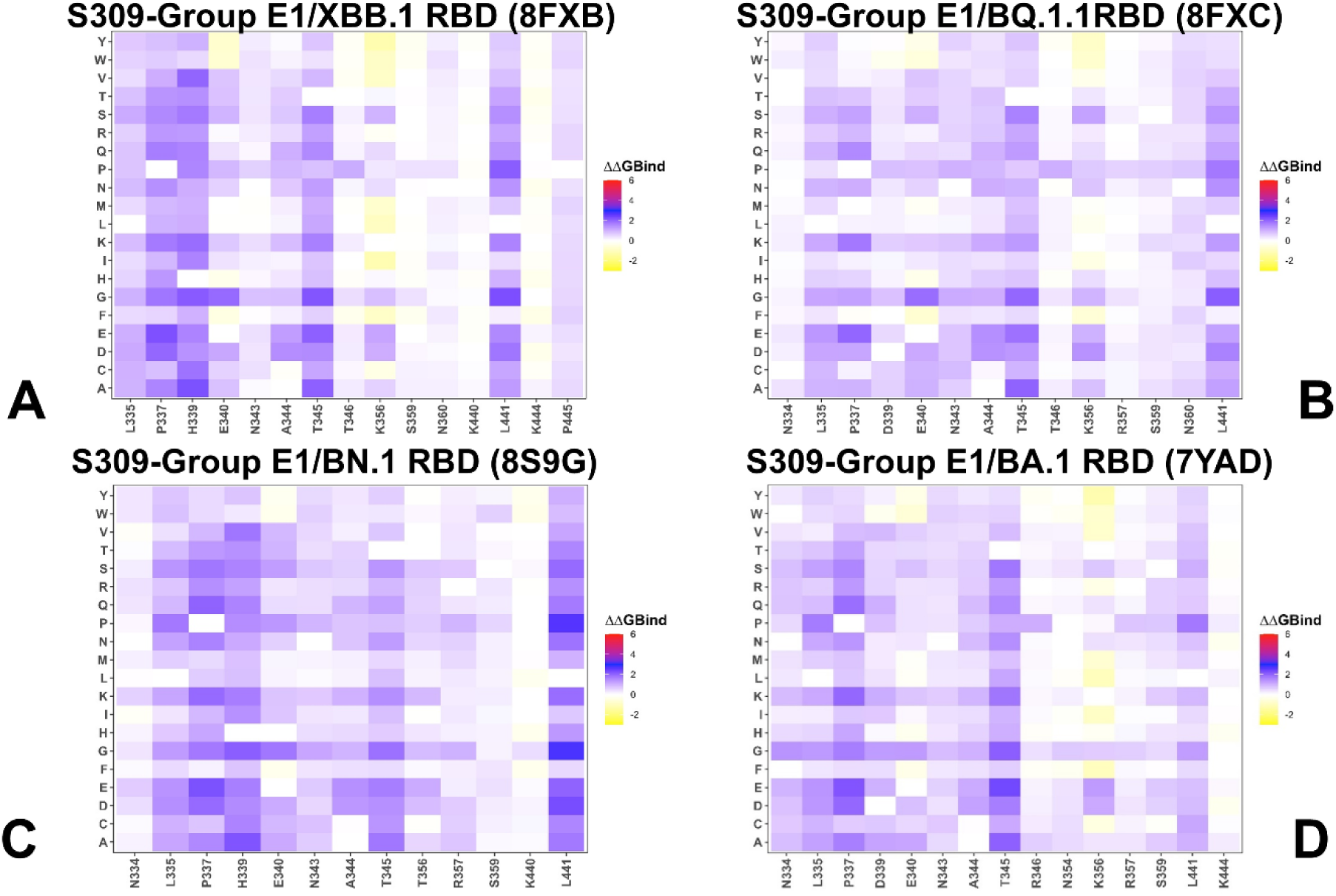
The ensemble-based mutational scanning of binding for the SARS-CoV-2 S-RBD complexes with S309 antibody. The mutational scanning heatmaps for the binding epitope residues in the S-RBD complexes with XBB.1 RBD, pdb I 8FXB (A), BQ.1.1 RBD, pdb id 8FXC (B), BN.1 RBD, pdb id 89GG (C) and BA.1 RBD, pdb id 7YAD (D). The binding energy hotspots correspond to residues with high mutational sensitivity. The heatmaps show the computed binding free energy changes for 20 single mutations on the sites of variants. The squares on the heatmap are colored using a 3-colored scale blue-white-yellow, with yellow indicating the largest unfavorable effect on stability. The standard errors of the mean for binding free energy changes were based on a 500 samples from MD trajectory are within 0.08-0.15 kcal/mol.

Specifically, N334 forms hydrogen bonds with the S309 paratope, stabilizing the interaction. Substitutions at N334 could disrupt these hydrogen bonds, reducing the binding affinity of S309. This makes N334 a critical residue for maintaining high-affinity interaction. L335 contributes to hydrophobic contacts with the S309 antibody. Hydrophobic interactions are essential for burying surface area at the antibody-RBD interface, ensuring tight binding. Mutations at L335 could introduce polar or charged residues, disrupting the hydrophobic core and destabilizing the interaction. P337 is located within the helical region and plays a structural role in maintaining the conformation of the epitope. Proline residues often induce kinks or turns in secondary structures, which can influence the accessibility of the epitope. Substitutions at P337 could alter the local conformation of the helix, potentially affecting the orientation of nearby residues and impairing S309 binding.

Our results are consistent with the experiments that showed S309 binding is largely maintained against Omicron variants except BA.2.75.2, CA.3, CH.1.1, BA.2.86, and JN.1 variants. CH.1.1 acquired the R346T and F486S mutations present in BA.2.75.2 as well as harbored the additional K444T and L452R. CA.3.1 variant acquired R346T, F486S, K444M and L452R mutations [103]. This study showed that R346T could confer strong neutralization resistance to S309-like antibodies in CH.1.1 and CA.3.1 [103]. A series of SARS-CoV-2 variants with mutations at the L455, F456, and R346 positions include the "SLip" variant, which carries L455S along with an additional F456L mutation. More recently, the "FLiRT" variant has appeared, featuring an additional R346T mutation on the SLip backbone. Studies have shown that the SLip and FLiRT subvariants of JN.1 exhibited a complete escape of neutralization by S309 [104]. Residues D339 and R346 lie within the epitope region of S309. Mutations at these position are present in lineages FLiRT and KP.2, which could enhance viral evasion from antibody neutralization. Several studies showed S309 maintained efficacy against Omicron variants including BA.2.87.1 with the exception of CH.1.1, CA.3.1, BA.2.75.2, and BA.2.86 [103-105]. BA.2.87.1 variant features N417T, K444N, V445G, L452M, K478T, N481K, R493Q [105] and can be effectively neutralized by S309. Across all four S309-RBD complexes, T345, R346, and L441 positions correspond to mutational hotspots and variations in these sites, particularly R346T convergent mutation can render resistance to S309 binding (Figure 5, Supporting Information, Datasets S1-S4). Mutational scanning also revealed P337 position as an important hotspot for S309 binding (Figure 5, Supporting Information, Datasets S1-S4). This is consistent with reported data that any substitution of the helix breaker P337 at resulted in a complete S309 escape [106].

Mutational scanning of the RBD in complexes with the S304 antibody across four different structures revealed consistent hotspots that play critical roles in stabilizing the interaction. These hotspots include residues Y369, F374, F377, C379, Y380, G381, V382, S383, P384, T385, F390, and L392. Among these, S383, P384, and T385 emerged as the most significant energy hotspots (Figure 6, Supporting Information, Datasets S5-S8). T385 serves as the dominant hotspot for S304 binding, acting as a central energetic hub and its critical role is underscored by experimental data showing that even subtle mutations (e.g., T385K/D/E/R) disrupt binding affinity [27,39,40]. The high sensitivity of this residue highlights its importance in maintaining the structural integrity of the epitope and suggests that it is a key target for immune evasion by the virus. S383 and P384 contribute to the dense network of interactions within the epitope. P384, in particular, acts as a structural anchor due to its rigid proline ring, which helps maintain the local conformation necessary for optimal binding. Together, these residues ensure tight packing at the interface, providing additional stability to the complex. Y380 participates in hydrophobic interactions, forming part of the core of the epitope. The identification of the Y380Q mutation in the B.1.91 lineage suggests that substitutions at this position may reduce hydrophobic packing, potentially destabilizing the interaction between S304 and the RBD. The RBD novel mutation (Y380Q) was found in one sample occurring simultaneously with C379W and V395A, and the B.1.91 lineage in the spike protein. The Y380Q and C379W may interfere with the binding of neutralizing antibody S304 [107]. K386 plays a crucial role in electrostatic stabilization, working synergistically with T385 to enhance binding affinity. The deleterious effects of mutations like K386D/R indicate that disruptions to this electrostatic network can significantly impair S304 binding [27,39,40]. The identification of key hotspots T385, S383, and P384 provides valuable insights into the mechanisms of immune escape. These findings align with DMS data [27], confirming the robustness of the computational predictions. The consistency of the results across multiple structures emphasizes the validity of the identified hotspots and their relevance to understanding viral evolution and immune evasion.

**Figure 6.**
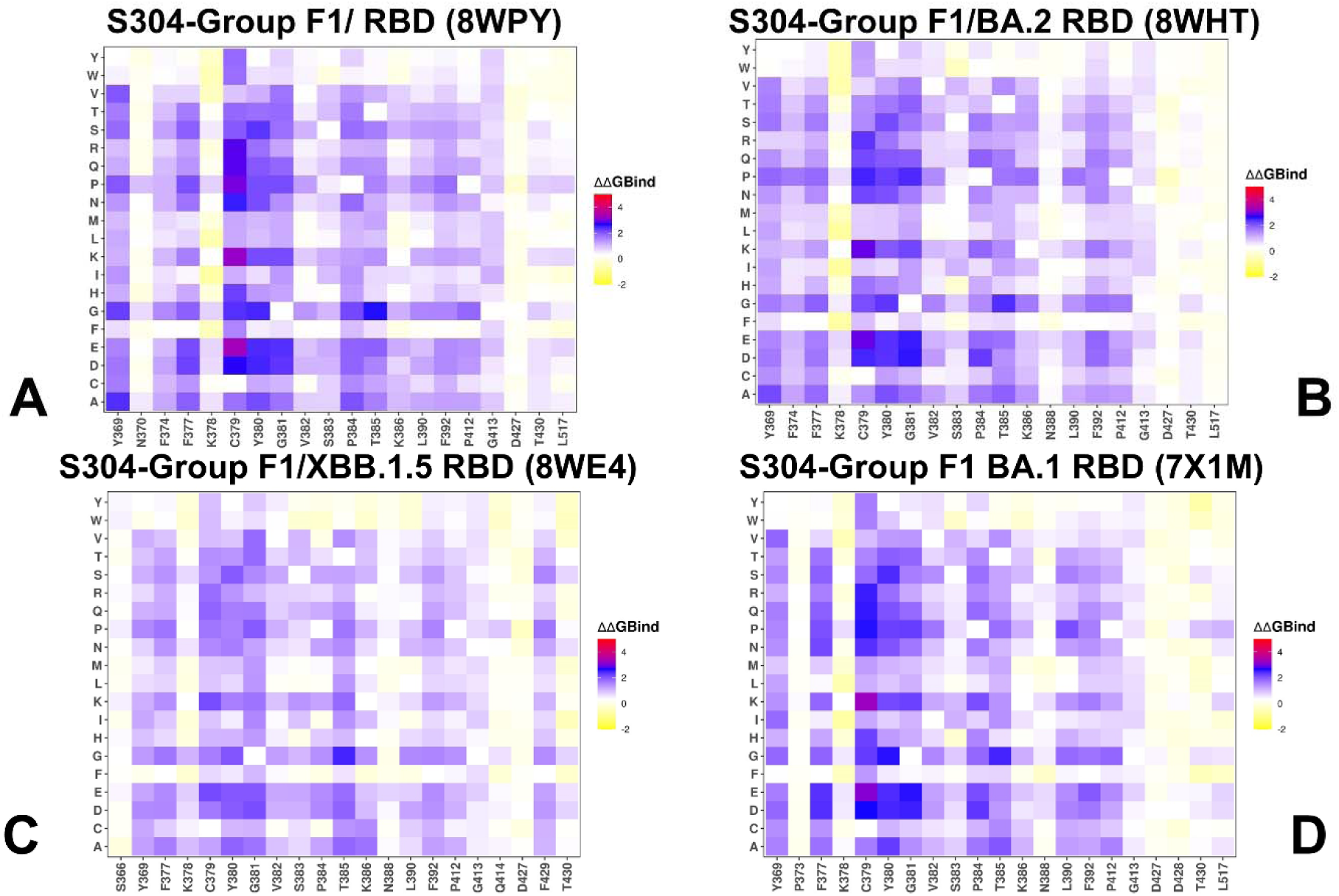
The ensemble-based mutational scanning of binding for the SARS-CoV-2 S-RBD complexes with S304 antibody. The mutational scanning heatmaps for the binding epitope residues in the S304 complexes with Wu-WT RBD, pdb id 8WPY (A), BA.2 RBD, pdb id 8HWT (B), XBB.1.5 RBD, pdb id 8WE4 (C) and BA.1 RBD, pdb id 7X1M (D). The binding energy hotspots correspond to residues with high mutational sensitivity. The heatmaps show the computed binding free energy changes for 20 single mutations on the sites of variants. The squares on the heatmap are colored using a 3-colored scale blue-white-yellow, with yellow indicating the largest unfavorable effect on stability. The standard errors of the mean for binding free energy changes were based on 500 selected samples from MD trajectory are within 0.12 kcal/mol.

Mutational scanning of the RBD residues in complexes with CYFN-1006.1 and CYFN-1006.2 antibodies (Figure 7A,B, Supporting Information, Datasets S9,S10) consistently unveiled key energetic hotspots at positions A344, T345, T346, F347, L441 and P499. Importantly, there are no notable Omicron variants that carry mutations in positions A344, T345, F347, L441 and P499. However, a newly discovered point R346X, primarily R346T, is connected with surge in SARS-CoV-2 infections as R346T mutation is predominantly expressed in many Omicron subvariants. Generally, the three notable amino acid substitutions at the R346 position of the spike RBD are R346K, R346T, and R346I. Most of the offspring descended from the Omicron sublineages, including BJ.1, BR.3, and BA.2.75.5, show a strong predominance of R346T [108]. R346K is in B1.62.1 and BA.1.1 [109]. Notably, we used ensemble from simulations of XBB.1.16 and EG.5.1 RBD in complexes with antibodies CYFN1006-1/2 in which positions T345 and T346 are present. It appears that mutations in T346 position can moderately reduce binding significantly and R346T is present in these variants as well as in KP.2 but not in JN.1 and KP.3 (Figure 7A,B, Supporting Information, Datasets S9,S10). CYFN1006-1 was tested against B.1.1.7, B.1.351, P.1, B.1.617.2, BA.1 and Variants of Interest (VOIs) (B.1.525, B.1.621, C.37) as well as various XBB subvariants, along with recently identified and currently circulating variants such as JN.1, KP.2, KP.3, KP.3.1.1 and XEC [41].

**Figure 7.**
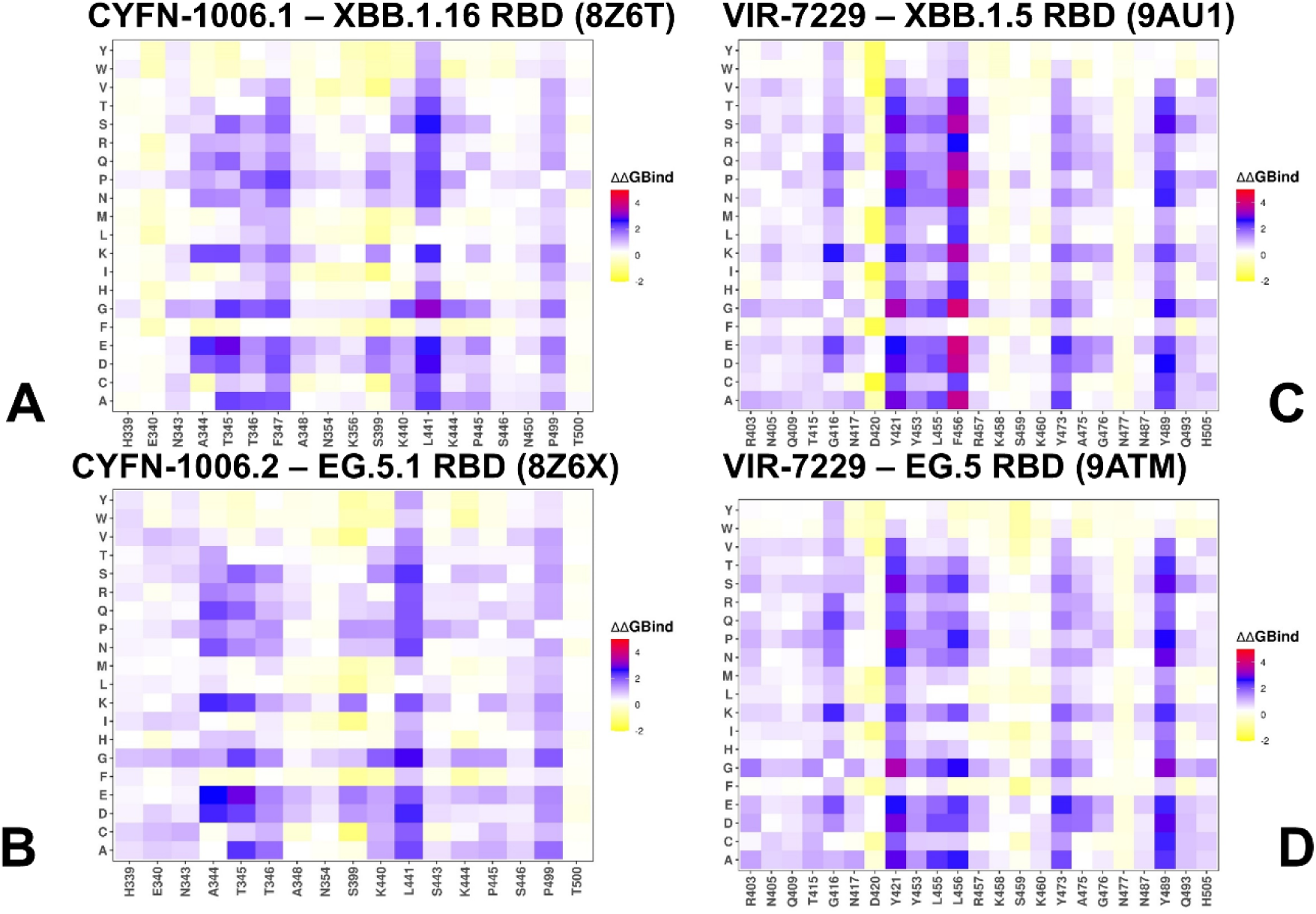
The ensemble-based mutational scanning of binding for the SARS-CoV-2 S-RBD complexes with CYFN-1006 (A,B) and VIR-7229 (C,D). The mutational scanning heatmaps for the binding epitope residues in the CYFN-1006.1 complex with XBB.1.16 RBD, pdb id 8Z6T (A), CYN-1006.2 complex with EG.5.1 RBD, pdb id 8Z6X (B). The heatmaps for VIR-7229 complexes with XBB.1..5 RBD, pdb id 9AU1 (C) and EG.5 RBD, pdb id 9ATM (D). The binding energy hotspots correspond to residues with high mutational sensitivity. The heatmaps show the computed binding free energy changes for 20 single mutations on the sites of variants. The squares on the heatmap are colored using a 3-colored scale blue-white-yellow, with yellow indicating the largest unfavorable effect on stability. The standard errors of the mean for binding free energy changes were based on 500 selected samples from MD trajectory are within 0.21 kcal/mol.

Of particular interest are mutational heatmaps of VIR-7229 binding to XBB.1.5 RBD (Figure 7C, Supporting Information, Dataset S11) and EG.5 RBD (Figure 7D, Supporting Information, Dataset S12). These maps are highly similar and point to mutational hotspots at positions G416, Y421, Y453, L455, F456/L456, Y473 and Y489. Hence, despite large binding interface, a hydrophobic group of highly conserved central residues are associated with VIR-7229 binding. With a notable exception of L455 and F456 positions other residues have a high mutational barrier and typically not observed in Omicron variants. KP.2 that carries mutations R346T, F456L, and KP.3 that features mutations R346T, L455S, F456L, Q493E, share the F456L mutation and KP.3 has emerged as the most immune-evasive and fastest-growing JN.1 sublineage, largely due to the F456L mutation, which plays a critical role in the antibody escape [63]. Mutational heatmaps showed that F456 can be more favorable for binding, while L456 mutations are somewhat better tolerated but still highlight the binding hotspot for both F456 and L456 (Figure 7C,D, Supporting Information, Datasets S11,S12).

Our results are in line with the DMS experiments that exhaustively mapped its escape profile of the Wuhan-Hu-1, BA.2, BQ.1.1, XBB.1.5, EG.5, and BA.2.86 yeast-displayed RBDs. VIR-7229 featured a remarkably narrow escape profile in all backgrounds with only some mutations at F456 causing escape [42]. A direct comparison with mutational escape in the XBB.1.5 background featured in the structure of VIR-7229 (Figure 7C, Supporting Information, Dataset S11), the mutations causing significant escape are F486K/E/P [42]. Our results showed that largest destabilization free energies for mutations of F456A (ΔΔG = 3.75 kcal/mol), F456D (ΔΔG = 3.74 kcal/mol), F456K (ΔΔG = 3.49 kcal/mol), F456E (ΔΔG = 3.94 kcal/mol) and F486P (ΔΔG = 3.82 kcal/mol) (Figure 7C, Supporting Information, Dataset S11). More tolerant changes are evident for VIR-7229 binding with EG.5 RBD which featured F456L modification (Figure 7D, Supporting Information, Dataset S12). Interestingly, mutations L456P and L456S emerged as highly destabilizing which is consistent with the experiments. To summarize, mutational scanning identified specific residues that are crucial for antibody binding, revealing particularly commonalities and specifics of mutational resistance patterns. We found that certain mutations, particularly R346T and F456L, are consistently associated with reduced antibody efficacy across multiple variants. These mutations highlight the evolutionary pressure on the virus to evade immune responses. Different antibodies exhibit unique escape profiles, with some showing broader efficacy and others being more susceptible to specific mutations. For instance, S309 shows resilience against many variants but fails against those with R346T and F486S mutations. Despite the presence of numerous mutations, certain highly conserved residues maintain their importance in antibody binding, suggesting that they may be less prone to mutational escape. Some conserved residues directly participate in critical interactions with antibodies. For instance, N334 forms hydrogen bonds with the S309 paratope, stabilizing the interaction. Substitutions at this position could disrupt these bonds, reducing binding affinity. F456 in the VIR-7229 epitope is part of a hydrophobic cluster that is central to the binding interface. Although mutations at F456 can destabilize binding, F456L variant can bind to VIR-7229 through mutual accommodation of the binding partners. In general, these residues are functionally indispensable for maintaining high-affinity interactions, making them less likely to tolerate mutations. Conserved residues represent a "weak spot" in the virus’s ability to evolve resistance, offering a stable target for intervention. Understanding the conservation patterns of key residues can help predict which mutations are likely to emerge and their potential impact on immune evasion. The prevalence of R346T mutations in Omicron subvariants underscores the evolutionary advantage of this mutation, but the rarity of mutations at positions like F456 highlights their critical role in maintaining viral fitness. These residues offer stability to the viral protein, making them less prone to mutational escape. Furthermore, understanding the evolutionary constraints on these residues can inform strategies to anticipate and counteract future mutations.

### MM-GBSA Analysis of the Binding Affinities

The experimental and computational studies suggested that the cross-neutralization Ab activity against Omicron variants may be driven by balance and tradeoff of multiple energetic factors and interaction contributions of the evolving escape hotspots involved in antigenic drift and convergent evolution. However, the dynamic and energetic details quantifying the balance and contribution of these factors, particularly the balancing nature of specific interactions formed by Abs with the epitope hotspot residues remains scarcely characterized. Here, using the conformational equilibrium ensembles obtained MD simulations we computed the binding free energies for the RBD-Ab complexes using the MM-GBSA method [105-108]. In this analysis, we focused on the binding free energy decomposition and examination of the energetic contributions of individual RBD epitope residues. Through this analysis, we determined the binding hotspots for Ab binding and quantified the role of the van der Waals and electrostatic interactions in the binding mechanism. In the MM-GBSA calculations, we examine whether the binding affinities and contributions of the major binding hotspots are largely determined by the van der Waals or electrostatic interactions and whether positions of immune escape can be associated with the binding hotspots where different energetic contributions act synergistically leading to significant loss of binding upon mutations.

MM-GBSA analysis of S309 antibody complexes and residue decomposition of the total energy (Supporting Information, Tables S2-S5) revealed strong and consistent binding hotspots for T345(ΔG = -4.72 kcal/mol), K356 ((ΔG = -3.29 kcal/mol), L441 (ΔG = -2.83 kcal/mol) N343 (ΔG = -3.03 kcal/mol), A344 (ΔG = -1.72 kcal/mol), and R346 positions (ΔG = -1.69 kcal/mol) (Figure 8A, Supporting Information, Tables S2-S5). This is consistent with mutational scanning computations that identified these residues as critical for binding S309 antibody.

**Figure 8.**
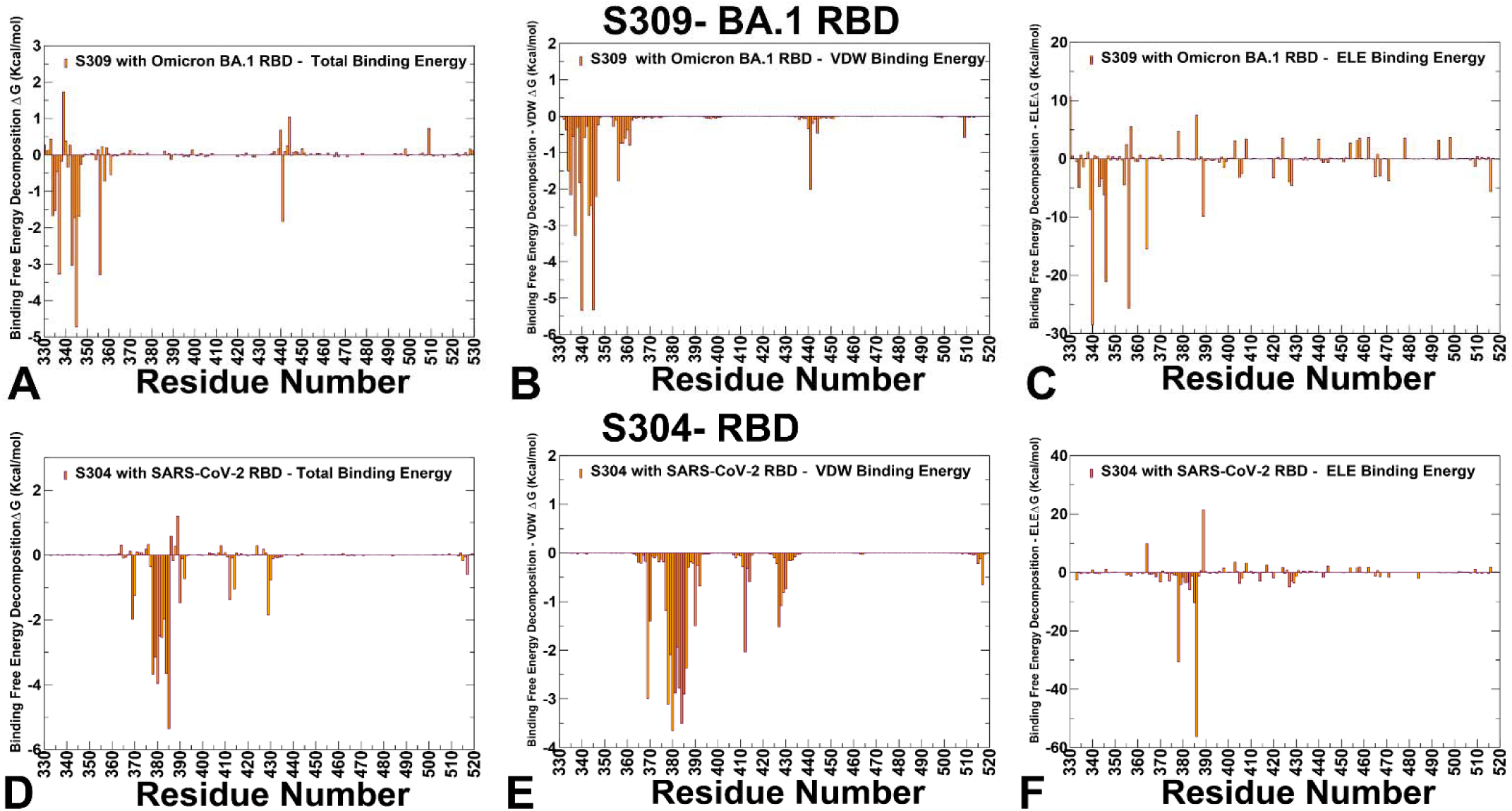
The residue-based decomposition of the binding MM-GBSA energies for S309 and S304 binding. The residue-based decomposition for S309-RBD binding shows the total binding energy (A), van der Walls contribution to the total MM-GBSA binding energy (B) and the electrostatic contribution to the total binding free energy (C). MM-GBSA contributions are evaluated using 1,000 samples from the equilibrium MD simulations of S309 complex with BA.1 RBD complex, pdb id 7YAD. The residue-based decomposition for S304-RBD binding shows the total binding energy (D), van der Walls contribution to the total MM-GBSA binding energy (E) and the electrostatic contribution to the total binding free energy (F). MM-GBSA contributions are evaluated using 1,000 samples from the equilibrium MD simulations of S304 complex with Wu-WT RBD, pdb id 8WPY.

The energy decomposition showed that the strongest van der Waals interactions are provided by T345, P337, A344, L335 and L441 RBD residues (Figure 8B, Supporting Information, Tables S2-S5). T345 and P337 are key hotspots that are driven by favorable hydrophobic contacts. The largest electrostatic interactions are formed by residues E340, K356, R346 and D389 (Figure 8C, Supporting Information, Tables S2-S5). The results indicate that the emergence of T345, K356 and R346 positions as key hotspots may arise from synergistic favorable van der Waals and electrostatic interactions whereas P337 and L441 positions are largely involved in packing interactions. These findings can explain why R346S in combination with P337L enhanced resistance to S309 [34]. This is consistent with reported data that any substitution of the helix breaker P337 at resulted in a complete S309 escape [106]. R346K in VOC Omicron BA.1.1 reduced sensitivity to S309 [35]. The experiments showed that neutralization activity against BQ, CH, XBB, showed reduced S309 binding and complete loss of binding for JN.1, KP.2, KP.3, and XEC variants all of which share K356T mutation [103-105]. Our results showed that K356 is the second most dominant hotspot of S309 binding delivering the most favorable electrostatic contribution which may explain resistance of JN.1, KP.2 and KP.3 variants to S309. Binding free energy analysis of S304 binding revealed S383, T385, Y380, K378, P384, C379, V382 as most significant energy hotspots with T385 (ΔG = -5.36 kcal/mol) emerging as a dominant energetic center (Figure 8D, Supporting Information, Table S6). While van der Waals interactions are strongest for Y380, P384, Y369 and T385, the electrostatics favors K386, K378 and also T385 (Figure 8E, Supporting Information, Table S6). Strikingly, the experimental data on escape hotspots singled out S383, T385 and K386 sites [27,39,40]. Interestingly, both van der Waals, and electrostatic interactions work synergistically to yield fa favorable binding or T385 and K386 (Figure 8F, Supporting Information, Table S6). These findings are also consistent with mutational scanning computations yielding correct immune hotspots and reproducing DMS data [27]. S383, P384, and T385 emerged as the most significant energy hotspots based on escape maps and binding free energy analysis. T385 consistently emerged as the dominant energetic center with a binding free energy contribution of ΔG = -5.36 kcal/mol. It is stabilized by both strong van der Waals interactions and electrostatic forces, making it indispensable for S304 binding. S383 and P384 contribute significantly to the stability of the complex through van der Waals interactions, with P384 playing a structural role due to its rigidity, likely influencing the local conformation of the epitope.

To conclude we highlight some important patterns and trends of S309 and S304 binding mechanisms. The binding of S309 is predominantly driven by a combination of hydrophobic packing and electrostatic interactions. Key residues like T345, P337, L441, and R346 play dual roles in stabilizing the complex through van der Waals and electrostatic forces. The synergy between these interactions explains why mutations at R346 (e.g., R346T/K/S) confer resistance to S309. The residues involved in S309 binding, such as T345, P337 and R346, are structurally and functionally constrained, making them less prone to random mutations. However, when mutations do occur (e.g., R346T), they can have a profound impact on binding affinity. S304 binding is more localized around specific residues, with T385 emerging as the dominant energetic hotspot. Both van der Waals, and electrostatic interactions contribute synergistically to stabilizing the complex. Mutations at T385 and K386 are highly deleterious, indicating their critical roles in maintaining binding affinity. Unlike S309, which relies on multiple residues spread across the epitope, S304 leverages contribution of fewer critical residues. The immune escape profile of S309 is linked to mutations at R346 and P337 which disrupt the balance of van der Waals and electrostatic interactions. These residues are conserved or minimally mutated in many variants, except in specific subvariants like BA.2.75.2, CH.1.1, and BA.2.86 where mutations like R346T confer strong resistance. The presence of convergent mutations like R346T highlights the evolutionary pressure on the virus to evade S309 neutralization while maintaining fitness. For S304, immune escape is primarily associated with mutations at T385 and K386, which are energetically dominant hotspots. The experimental data align with computational predictions, showing that mutations like T385K/D/E/R and K386D/R significantly impair binding [27]. The residues involved in S304 binding, particularly T385 and K386, are also under evolutionary constraints due to their critical roles in antibody binding. However, the localized nature of S304’s binding energy makes it more susceptible to escape mutations at these specific sites. This localized dependence contrasts with S309’s broader binding network, which may offer some resilience against single-point mutations.

CYFN1006-1 largely overlap with S309 situated on the outer surface of RBD near the RBM region with the epitopes comprising N343, T345, R346, K356 in S309 epitope, forming numerous hydrogen bonds and displaying high conservation. The binding affinity of CYFN-1006.1 and CYFN-1006.2 antibodies is driven primarily by N343, A344, T345, K440, T346, P445, S446, A344, L441 (Figure 9, Supporting Information, Tables S7,S8). The van der Waals interactions are most favorable for K440, T345 and L441 while the electrostatics favors K440, K444. D364. T365 and D389 (Figure 9B,E, Supporting Information, Tables S7,S8). Importantly, K356 position does not provide significant contribution to binding of these antibodies which may provide enhanced neutralization potency against JN.1, KP.2 and KP.3 variants sharing K356T mutaion. N343, A344, and T345 play an important role in stabilizing CYFN1006-1 binding to RBD and are highly conserved in various mutant strains. K440, L441, P445, and P499 of RBD are embedded in the hydrophobic patch formed by CDRH1 (Y36, Y37, W38) and CDRH3 (W112) to further stabilize the CYFN1006-1 binding to RBD.

**Figure 9.**
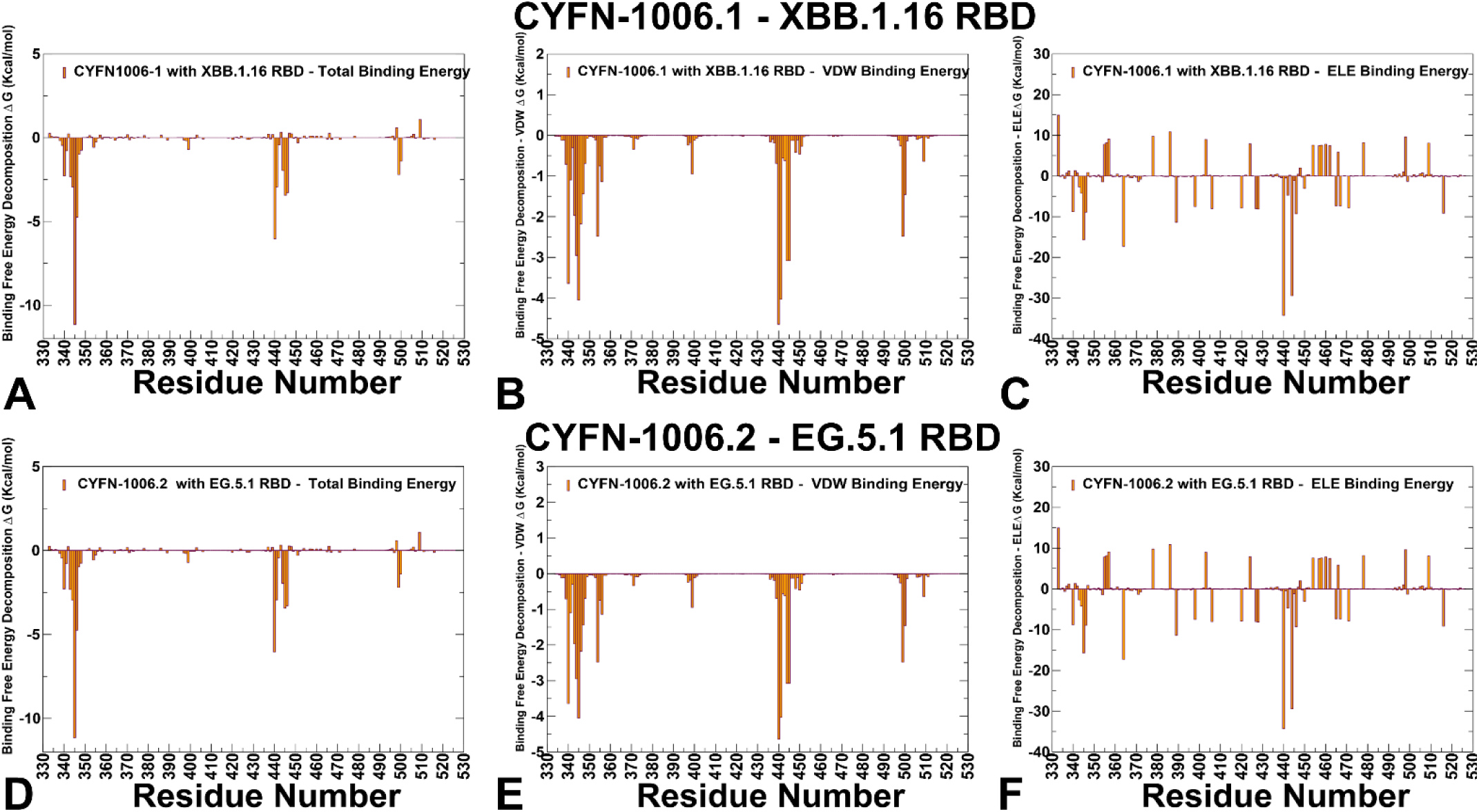
The residue-based decomposition of the binding MM-GBSA energies for CYFN-1006 binding. The residue-based decomposition for CYFN-1006.1 -XBB.1.16 RBD binding shows the total binding energy (A), van der Walls contribution to the total MM-GBSA binding energy (B) and the electrostatic contribution to the total binding free energy (C). MM-GBSA contributions are evaluated using 1,000 samples from the equilibrium MD simulations of CYFN-1006.1 complex with XBB.1.16 RBD complex, pdb id 8Z6T. The residue-based decomposition for CYFN-1006.2 -EG.5.1 RBD binding shows the total binding energy (D), van der Walls contribution to the total MM-GBSA binding energy (E) and the electrostatic contribution to the total binding free energy (F). MM-GBSA contributions are evaluated using 1,000 samples from the equilibrium MD simulations of C YFN-1006.2 complex with EG.5.1 RBD, pdb id 8Z6X.

A direct comparison with mutational escape in the XBB.1.5 background featured in the structure of VIR-7229 the mutations causing significant escape are F486K/E/P [42]. This may explain that VIR-7229 interactions can tolerate epitope variability exhibiting high barrier for the emergence of resistance, partly attributed to its high binding affinity [43]. In particular, VIR-7229 showed neutralization efficacy against KP.2 that carries mutations R346T, F456L, and KP.3 that features mutations R346T, L455S, F456L, Q493E. Although R403 and F456L mutations are present in KP.2 and KP.3 variants and arguably can reduce neutralization potency of VIR-7229, their effect is relatively forgiving. Out of the 11 mutated residues in BA.2.86 RBD relative to XBB.1.5, only R403K is found in the VIR-7229 epitope. Our results suggest that R403K preserve electrostatic interactions with the VIR-7229 light-chain N33 and D52 side chains (Figure 10C,F, Supporting Information, Tables S9,S10). JN.1 variant harbors the immune-evasive L455S mutation and compatible with the VIR-7229 paratope interface due to the small size of the introduced serine side chain. Consistent with these results, mutational scanning of the RBD residues in complexes with CYFN-1006.1 and CYFN-1006.2 antibodies (Figure 7A,B) showed that key energetic hotspots at positions A344, T345, T346, F347, L441 and P499. Similarly to S309, these antibodies target T345 and T346 sites as dominant hotspots. Even though binding epitope of CYFN-1006-1 includes 17 RBD residues, the conserved residue T345 along with K440 and T346 anchor binding affinity of these potent neutralizing antibodies. However, according to the experiments, neutralization activity against BQ, CH, XBB, JN.1, KP.2, KP.3, and XEC variants showed reduced binding or complete loss of binding for S309, while CYFN1006-1 demonstrated consistent neutralization of all tested SARS-CoV-2 variants with unaffected potency.

**Figure 10.**
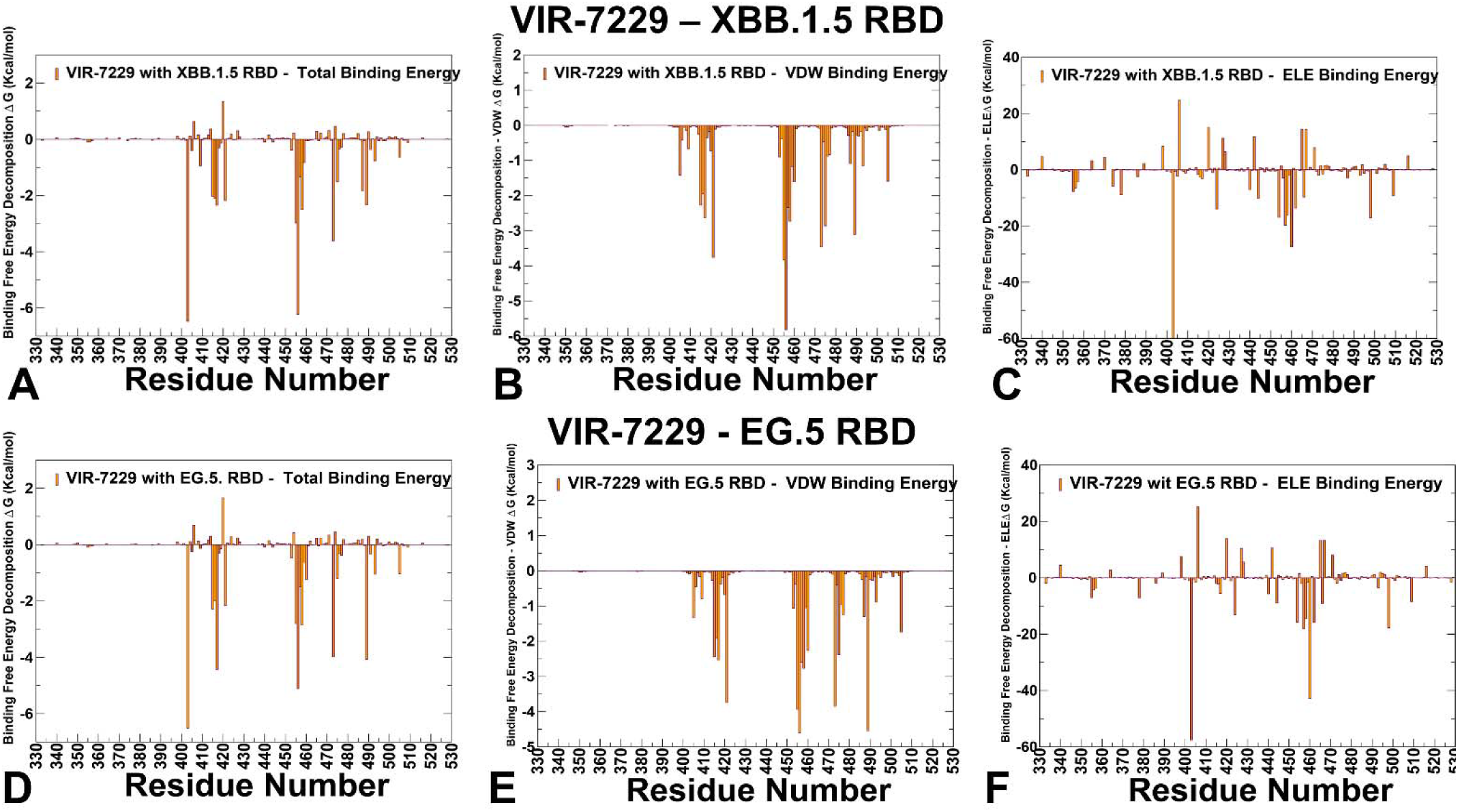
The residue-based decomposition of the binding MM-GBSA energies for VIR-7229 binding. The residue-based decomposition for VIR-7229 -XBB.1.5 RBD binding shows the total binding energy (A), van der Walls contribution to the total MM-GBSA binding energy (B) and the electrostatic contribution to the total binding free energy (C). MM-GBSA contributions are evaluated using 1,000 samples from the equilibrium MD simulations of VIR-7229complex with XBB.1.5 RBD complex, pdb id 9AU1. The residue-based decomposition for VIR-7229 - EG.5 RBD binding shows the total binding energy (D), van der Walls contribution to the total MM-GBSA binding energy (E) and the electrostatic contribution to the total binding free energy (F). MM-GBSA contributions are evaluated using 1,000 samples from the equilibrium MD simulations of VIR-7229 complex with EG.5 RBD, pdb id 9ATM.

Binding affinity of VIR-7229 to XBB.1.5 RBD showed hotspot in positions R403 (ΔG = -6.47 kcal/mol), F456 (ΔG = -6.23 kcal/mol), Y473 (ΔG = -3.61 kcal/mol) and L455 (ΔG = -2.97 kcal/mol) (Figure 10A, Supporting Information, Table S9). The most favorable van der Waals contacts are made by F456 (ΔG = -5.82 kcal/mol), L455, Y421, Y473 and Y489 residue, while the electrostatics is most favorable for R403, K460, R457 and R498 (Figure 10B, Supporting Information, Table S9). Interestingly, VIR-7229 binding to EG.5 variant with F456L mutation preserves the ranking of major hotspots R403, L456, 417, Y489, Y473, K458 and L455 (Figure 10D, Supporting Information, Table S10) as L456 makes favorable contacts allowing for optimal complementarity with L456 (ΔG = -5.11 kcal/mol). The van der Waals interactions for L456 (ΔG = -4.6 kcal/mol) are only moderately weaker than for F456, but the overall binding contribution of mutated F456L continues to be highly favorable (Figure 10E, Supporting Information, Table S10). A direct comparison with mutational escape in the XBB.1.5 background featured in the structure of VIR-7229 the mutations causing significant escape are F486K/E/P [42]. This may explain that VIR-7229 interactions can tolerate epitope variability exhibiting high barrier for the emergence of resistance, partly attributed to its high binding affinity [43]. In particular, VIR-7229 showed neutralization efficacy against KP.2 that carries mutations R346T, F456L, and KP.3 that features mutations R346T, L455S, F456L, Q493E. Although R403 and F456L mutations are present in KP.2 and KP.3 variants and arguably can reduce neutralization potency of VIR-7229, their effect is relatively forgiving. Out of the 11 mutated residues in BA.2.86 RBD relative to XBB.1.5, only R403K is found in the VIR-7229 epitope. Our results suggest that R403K preserve electrostatic interactions with the VIR-7229 light-chain N33 and D52 side chains (Figure 10C,F, Supporting Information, Tables S9,S10). JN.1 variant harbors the immune-evasive L455S mutation and compatible with the VIR-7229 paratope interface due to the small size of the introduced serine side chain.

In summary, the binding affinity of CYFN1006 is anchored by highly conserved residues such as T345, K440 and T346 which are critical for stabilizing the interaction through both van der Waals and electrostatic forces. Notably, K356 does not significantly contribute to binding, which may explain CYFN1006’s enhanced neutralization potency against variants carrying the K356T mutation. VIR-7229 targets a distinct set of residues, with R403, F456, Y473, and L455 emerging as dominant energetic hotspots. The antibody relies on a combination of hydrophobic and electrostatic interactions, with F456 playing a central role in hydrophobic packing and R403 contributing strongly to electrostatic stabilization. While residues F456 and L455 are critical for binding, VIR-7229’s adaptability to mutations at these positions suggests a broader tolerance for evolutionary changes. The antibody’s reliance on a hydrophobic core of highly conserved residues (Y473, Y489) ensures stability, while its flexibility allows it to accommodate substitutions.

## Discussion

The results presented in this study provide a comprehensive understanding of the molecular mechanisms underlying the interactions between SARS-CoV-2 spike protein receptor-binding domain (RBD) and neutralizing antibodies, including S309, S304, CYFN1006, and VIR-7229. These findings reveal key insights into the structural, energetic, and mutational determinants of antibody binding, shedding light on the virus’s evolutionary strategies to evade immune responses and offering guidance for the design of therapeutics and vaccines. The analysis highlights that different antibodies employ distinct binding mechanisms, targeting unique epitopes on the RBD with varying degrees of conservation and flexibility. S309 binds near the ACE2 interface, leveraging conserved residues such as T345, P337, and R346 to stabilize its interaction through synergistic van der Waals and electrostatic forces. The reliance on multiple residues across the epitope provides some resilience against single-point mutations but renders it vulnerable to convergent mutations like R346T, which are prevalent in immune-evasive variants. S304 focuses its binding energy on fewer but sensitive residues, particularly T385 and K386, which serve as dominant energetic hotspots. While this localized dependence allows for strong binding, it also makes S304 more susceptible to escape mutations at these specific sites.CYFN1006 exhibits a broader epitope comprising 17 RBD residues, with T345, K440, and T346 serving as critical anchors. Notably, CYFN1006’s reduced reliance on K356 enhances its efficacy against variants carrying the K356T mutation, underscoring its potential as a broad-spectrum therapeutic. VIR-7229 targets a structurally complex epitope within the receptor-binding motif (RBM), with R403, F456, Y473, and L455 emerging as dominant hotspots. Its adaptability to mutations like F456L and L455S highlights its resilience to epitope variability, making it a promising candidate for combating immune-evasive variants. These differences in binding mechanisms underscore the importance of targeting diverse epitopes to counteract the virus’s ability to evolve resistance. Broad-spectrum antibodies like CYFN1006 and VIR-7229, which maintain efficacy against multiple variants, represent valuable tools in this regard. A recurring theme in the results is the role of highly conserved residues in stabilizing antibody-RBD interactions. For example, T345 and P337 are critical for S309 binding, with substitutions at these positions significantly impairing affinity. Similarly, T385 and K386 are indispensable for S304 binding. N343, A344, and T345 anchor CYFN1006 binding, while F456 and Y473 play central roles in VIR-7229 binding. These conserved residues represent "weak spots" in the virus’s ability to evolve resistance, as mutations at these positions could destabilize the RBD structure or impair essential functions like ACE2 binding. The mutational scanning and MM-GBSA analyses reveal consistent patterns of immune evasion across variants, highlighting specific mutations that confer resistance to antibodies. We found that R346T mutation, prevalent in Omicron subvariants, consistently reduces the efficacy of S309 and other antibodies targeting this region. This underscores the evolutionary pressure on the virus to exploit mutations that disrupt antibody binding while maintaining fitness. Mutations at F456 and L455, critical for VIR-7229 binding, demonstrate varying degrees of tolerance. While F456L is accommodated without significant loss of binding, mutations like L455S highlight the delicate balance between viral adaptation and immune evasion. K356T mutation in JN.1, KP.2, and KP.3 variants severely impairs binding of S309 antibody but does not affect CYFN1006, illustrating how subtle differences in epitope recognition can influence antibody performance. These findings emphasize the need for a nuanced understanding of how specific mutations impact antibody binding, guiding the development of interventions that remain effective against evolving variants.

## Conclusions

This study provides critical insights into the molecular mechanisms governing the interactions between the SARS-CoV-2 spike RBD and neutralizing antibodies, including S309, S304, CYFN1006, and VIR-7229. By integrating advanced computational techniques such as molecular dynamics simulations, mutational scanning, and binding free energy calculations, we uncovered the structural, energetic, and evolutionary determinants of antibody-RBD binding. These findings reveal the intricate balance between viral immune evasion and the host immune response, offering a foundation for designing effective interventions against SARS-CoV-2 and its variants. The analysis highlights the diversity of binding mechanisms employed by different antibodies, each targeting unique epitopes with varying degrees of conservation and flexibility. Antibodies like S309 and CYFN1006 demonstrate broad-spectrum efficacy by leveraging highly conserved residues, making them less susceptible to mutations that disrupt binding. In contrast, antibodies such as S304 exhibit localized dependence on fewer residues, rendering them more vulnerable to escape mutations. VIR-7229 stands out for its adaptability to mutations, maintaining strong binding affinity even in the presence of key substitutions like F456L and L455S. This adaptability underscores its potential as a resilient therapeutic option against immune-evasive variants. A recurring theme in the results is the critical role of conserved residues in stabilizing antibody-RBD interactions. Residues such as T345, F456, and Y473 emerge as evolutionary "weak spots," where mutations could destabilize the RBD structure or impair essential functions like ACE2 binding. These residues represent stable targets for therapeutic intervention, as they are less prone to random mutations due to their structural and functional importance. Understanding the evolutionary constraints on these residues can help predict future mutational trends and guide the design of vaccines and therapeutics that remain effective against emerging variants. The study also underscores the virus’s remarkable ability to evolve resistance through specific mutations, such as R346T and F456L, which consistently reduce antibody efficacy across multiple variants. These mutations reflect the selective pressure on the virus to evade immune responses while maintaining fitness. Broad-spectrum antibodies like CYFN1006 and VIR-7229, which tolerate such mutations, provide resilience to epitope variability offers a high barrier to resistance, making them valuable tools in combating the rapidly evolving landscape of SARS-CoV-2. The findings emphasize the importance of targeting diverse epitopes and conserved residues to counteract viral evolution. By leveraging antibodies with complementary binding mechanisms and focusing on regions less prone to mutational escape, advanced strategies for synergistic inhibition may be improved to stay ahead of the virus’s evolutionary trajectory.

## Supplementary Materials

The following supporting information can be downloaded at: www.mdpi.com/xxx/s1, Datasets S1-S4 present the results of mutational scanning for different S309-RBD complexes. Datasets S5-S8 present the results of mutational scanning for different S304-RBD complexes. Datasets S9,S10 present the results of mutational scanning for RBD complexes with CYFN-100.1 and CYFN-1006.2 antibodies. Datasets S11, S12 present the results of mutational scanning for VIR-7229 binding with XBB.1.5 RBD and EG.5 RBD respectively. Table S1 highlights mutational landscape for Omicron variants explored in this study. Tables S2-S5 present the results of MM-GBSA computations for different S309-RBD complexes. Tables S6 presents the results of MM-GBSA computations for S304-RBD complex. Tables S7,S8 present the results of MM-GBSA computations for RBD complexes with CYFN-1006.1 and CYFN.1006-2 respectively. Tables S9,S10 present the results of MM-GBSA computations for VIR-7229 complexes with XBB.1.5 RBD and EG.5 RBD.

## Author Contributions

Conceptualization, G.V.; Methodology, M.A., V.P., B.F., G.V.; Software, M.A., V.P., B.F., G.V.; Validation, G.V.; Formal analysis, G.V., M.A., V.P.,B.F.; Investigation, G.V.; Resources, G.V., M.A.; G.V.; Data curation, M.A., G.C., G.V.; Writing— original draft preparation, G.V.; Writing—review and editing, G.V.; Visualization, .A.; G.V. Supervision G.V. Project administration, G.V.; Funding acquisition, G.V. All authors have read and agreed to the published version of the manuscript.

## Institutional Review Board Statement

Not applicable.

## Informed Consent Statement

Not applicable.

## Data Availability Statement

The original contributions presented in this study are included in the article/supplementary material. Crystal structures were obtained and downloaded from the Protein Data Bank (http://www.rcsb.org). All simulations were performed using NAMD 2.13 package that was obtained from website https://www.ks.uiuc.edu/Development/Download/. All simulations were performed using the all-atom additive CHARMM36 protein force field that can be obtained from http://mackerell.umaryland.edu/charmm_ff.shtml. The rendering of protein structures was done with interactive visualization program UCSF ChimeraX package (https://www.rbvi.ucsf.edu/chimerax/) and Pymol (https://pymol.org/2/). All mutational heatmaps were produced using the developed software that is freely available at https://alshahrani.shinyapps.io/HeatMapViewerApp/.

## Acknowledgments

The authors acknowledge support from Schmid College of Science and Technology at Chapman University for providing computing resources at the Keck Center for Science and Engineering.

