## Supplemental Tables S1-S10 and Datasets S1-S10 for "Exploring Diverse Binding Mechanisms of Broadly Neutralizing Antibodies S309, S304, CYFN-1006 and VIR-7229 Targeting SARS-CoV-2 Spike Omicron Variants: Integrative Computational Modeling Reveals Balance of Evolutionary and Dynamic Adaptability in Shaping Molecular Determinants of Immune Escape": TableS1.docx

**Table S1.** Mutational landscape of the Omicron variants.

| **Omicron Variant** | **Mutational landscape** |
| --- | --- |
| BA.1 | A67, T95I, G339D, S371L, S373P, S375F, K417N, N440K,  G446S, S477N, T478K, E484A, Q493R, G496S, Q498R,  N501Y, Y505H, T547K, D614G, H655Y, N679K, P681H, N764K, D796Y, N856K, Q954H, N969K, L981F |
| BA.2 | T19I, G142D, V213G, G339D, S371F, S373P, S375F, T376A, D405N, R408S, K417N, N440K, S477N, T478K, E484A, Q493R, Q498R, N501Y, Y505H, D614G, H655Y, N679K, P681H, N764K, D796Y, Q954H, N969K |
| XBB.1.5 | T19I, V83A, G142D, Del144, H146Q, Q183E, V213E, G252V, G339H, R346T, L368I, S371F, S373P, S375F, T376A, D405N, R408S, K417N, N440K, V445P, G446S,N460K, S477N, T478K, E484A, F486P, F490S, R493Q reversal, Q498R, N501Y, Y505H, D614G, H655Y, N679K, P681H, N764K, D796Y, Q954H, N969K |
| BA.2.86 | T19I, R21T, S50L, del69-70,V127F, delY144, F157S, R158G, delN211, L213I, L226F, H25N,A264D, I332V, D339H, K356T, R403K, V445H, G446, N450D, L452W, N460K, N481K, del V483, A484K, F486P, R493Q, E554K, A570V, P612S, I670V, H68R, D939F, P1143L |
| JN.1 | T19I, R21T, S50L, del69-70,V127F, delY144, F157S, R158G, delN211, L213I, L226F, H25N,A264D, I332V, D339H, K356T, R403K, V445H, G446, N450D, L452W, **L455S,** N460K, N481K, del V483, A484K, F486P, R493Q, E554K, A570V, P612S, I670V, H68R, D939F, P1143L |
| KP.2 | T19I, R21T, S50L, del69-70,V127F, delY144, F157S, R158G, delN211, L213I, L226F, H25N,A264D, I332V, D339H, **R346T**, K356T, R403K, V445H, G446, N450D, L452W, **L455S, F456L**, N460K, N481K, del V483, A484K, F486P, R493Q, E554K, A570V, P612S, I670V, H68R, D939F, **V1104L,** P1143L |
| KP.3 | T19I, R21T, S50L, del69-70,V127F, delY144, F157S, R158G, delN211, L213I, L226F, H25N,A264D, I332V, D339H, K356T, R403K, V445H, G446, N450D, L452W, **L455S, F456L**, N460K, N481K, del V483, A484K, F486P, **Q493E**, E554K, A570V, P612S, I670V, H68R, D939F, **V1104L,** P1143L |
